# Structural Basis of Ligand-Specific Orthosteric-Allosteric Coupling and Sensory Tuning in the Human Bitter Taste Receptor TAS2R14

**DOI:** 10.64898/2026.05.14.725165

**Authors:** Tao Zhou, Jiening Wang, Chulun Wang, Zhen Han, Yi Zhang, Zishao Ouyang, Sheng Ye, Shan Wu, Anna Qiao

## Abstract

The bitter taste receptor TAS2R14 recognizes hundreds of structurally diverse ligands, yet the mechanisms governing its broad promiscuity, graded agonist efficacy, and multi-site regulation remain unclear. Here, we report cryo-EM structures of TAS2R14–Gi_1_ complexes bound to 3,5-diiodosalicylic acid (DA), flufenamic acid (FA) and aristolochic acid (AA) at resolutions of 2.44 Å, 2.39 Å, and 2.69 Å, with a clear efficacy hierarchy of AA > FA > DA. Transmembrane helix 6 (TM6) acts as a core regulatory hub, with agonist binding triggering 12 Å lever-like TM6 rearrangements to reshape receptor interfaces and modulate G-protein coupling strength. We also identify cholesterol hemisuccinate (CHS) as a new TAS2R14 orthosteric agonist, selectively potentiating DA-mediated signaling potency by ∼5-fold via allosteric-orthosteric coupling without affecting FA or AA. Our findings show TM6 dynamics unify TAS2R14’s key functional features, advancing insights into bitter receptor activation and sterol regulation.

## Introduction

The mammalian bitter taste perception system, mediated by the TAS2R family of G protein-coupled receptors (GPCRs), constitutes a crucial first line of defense by enabling the detection of a vast array of potentially toxic compounds^1^. TAS2Rs canonically exert physiological function by releasing calcium (Ca^2+^) and reducing cyclic adenosine monophosphate (cAMP) through G protein gustducin (including a taste-selective Gα subunit α_gust_) signaling ^2-4^, or alternatively through Gα_i_ signaling ^5-7^. Beyond taste, TAS2Rs function as extraoral chemosensors. In the airways, they trigger protective reflexes like cough and modulate immune responses, while their dysregulation contributes to asthma and chronic obstructive pulmonary disease (COPD) pathogenesis^8-10^.

A fundamental and long-standing question in sensory biology and GPCR pharmacology is how a relatively small repertoire of approximately 25 human TAS2Rs can recognize an immense chemical space of bitter agonists with remarkable breadth and diversity. Among these receptors, TAS2R14 stands out as an extreme generalist and a potential therapeutic target for respiratory diseases^11,12^, documented to respond to 385 structurally distinct compounds, including numerous pharmaceuticals, toxins, and natural products ^13^. This broad ligand spectrum features a high proportion of molecules containing diverse hydrophobic and aromatic ring systems, posing a significant mechanistic puzzle: how does a single receptor accommodate such chemical heterogeneity while translating distinct structural inputs into graded signaling outputs? General principles of GPCR activation, often derived from studies of monoamine or peptide receptors, typically involve a defined orthosteric pocket and a more limited set of endogenous ligands. In stark contrast, bitter taste receptors like TAS2R14 must possess intrinsic structural plasticity to engage myriad exogenous, often xenobiotic, agonists.

Recent studies using cryo-electron microscopy (cryo-EM) have revealed the presence of an orthosteric binding pocket occupied by cholesterol (CHL) as well as an intracellular allosteric site ^14-16^, yet the conformational landscape that dictates ligand-specific efficacy profiles, bridges diverse agonist binding and G protein coupling, and mediates crosstalk between distinct binding sites remains poorly defined.

Here, we address these central questions through an integrative structural and functional study of TAS2R14. We present high-resolution cryo-electron microscopy (cryo-EM) structures of the TAS2R14–Gi complex bound to three representative bitter agonists— 3,5-diiodosalicylic acid (DA), flufenamic acid (FA), and aristolochic acid (AA)— which share a common carboxylate moiety but possess divergent aromatic cores and ligand efficacies. These structures, together with functional and computational analyses, uncover a complicated multi-site binding and allosteric regulatory mechanism in TAS2R14. We identify a conserved intracellular allosteric pocket, an orthosteric site recognized by cholesterol and its analog cholesterol hemisuccinate (CHS), and a secondary cytoplasmic subpocket specific to certain agonists, whose distinct binding profiles drive unique conformational rearrangements of TM6 to modulate receptor activation and G protein coupling. Our findings define an orthosteric–allosteric coupling mode that tunes bitter agonist responses, offering a structural basis for the broad ligand spectrum of TAS2R14 and new insights into GPCR plasticity and ligand-directed signaling.

### Cryo-EM Structures of TAS2R14–Gi_1_ with DA, FA, AA: Diverse Scaffolds and Differential Efficacies

TAS2R14 is known for recognizing a broad spectrum of agonists, with 385 documented ligands in bitterDB (https://bitterdb.agri.huji.ac.il/dbbitter). Over one-third of these ligands are characterized by diverse cyclic ring architectures (Fig. S1). The representative agonists DA, FA, and AA all possess a conserved carboxylate moiety essential for receptor binding, yet they differ substantially in their hydrophobic aromatic cores, with structural complexity ranging from a simple phenol group in DA to an intricate polyaromatic scaffold in AA (Fig. 1A). Calcium mobilization assays have shown an efficacy ranking (E_max_) of AA (136±9%) > FA (100±5%) > DA (53±3%), indicating that DA has significantly lower efficacy compared to FA and AA (Fig. 1B, Table S4).

**Fig. 1.**
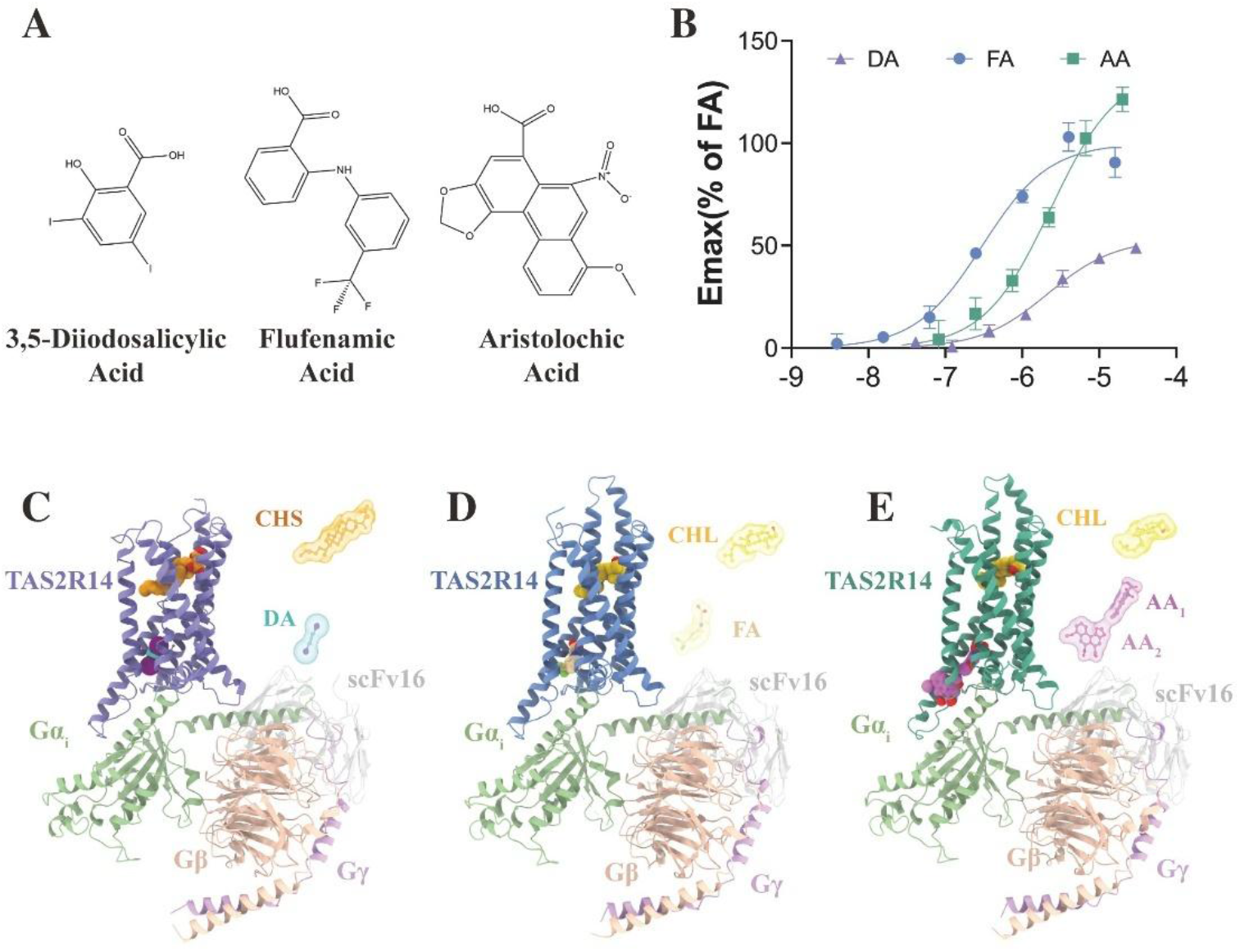
Chemically diverse agonists-dependent activation of the TAS2R14. (**A**) Chemical scaffolds of allosteric ligands for structural determination. (B) Ca^2+^ flux signal induced by DA, FA and AA. Data are means ± SEM of 4 biological replicates. (**C-E**) Cartoon representation of the DA-TAS2R14-G_i_ complex (C), FA-TAS2R14-G_i_ complex (D) and AA-TAS2R14-G_i_ complex (E). DA, FA and AA in TAS2R14-G_i_ complex structures are shown as cyan, beige and purple sticks, respectively. The DA-bound TAS2R14 is shown in violet, FA-bound TAS2R14 in royal blue, AA-bound TAS2R14 in green, G_αi_ subunit in lawn green, G_β_ subunit in salmon, G_γ_ subunit in light purple and scFv16 in gray.

To elucidate the structural basis for its broad ligand recognition and distinct ligand efficacies, we determined cryo-EM structures of the TAS2R14–Gi_1_ complex bound to DA, FA, and AA at 2.44 Å, 2.39 Å, and 2.69 Å resolution, respectively^17-22^ (Figure 1C-E, Fig. S2-3, Table S1). The overall structures of the three complexes are highly similar, with Cα RMSD values of 0.19 Å and 0.26 Å for DA-bound versus FA- and AA-bound states, respectively (Fig. S4).

### Ligand-specific interactions within the intracellular allosteric pocket of TAS2R14

Structural analysis revealed that DA, FA, and AA_1_ share a common allosteric pocket, while only the AA-bound complex accommodates a second ligand molecule, AA_2_ (Fig. 2A-E).

**Fig. 2.**
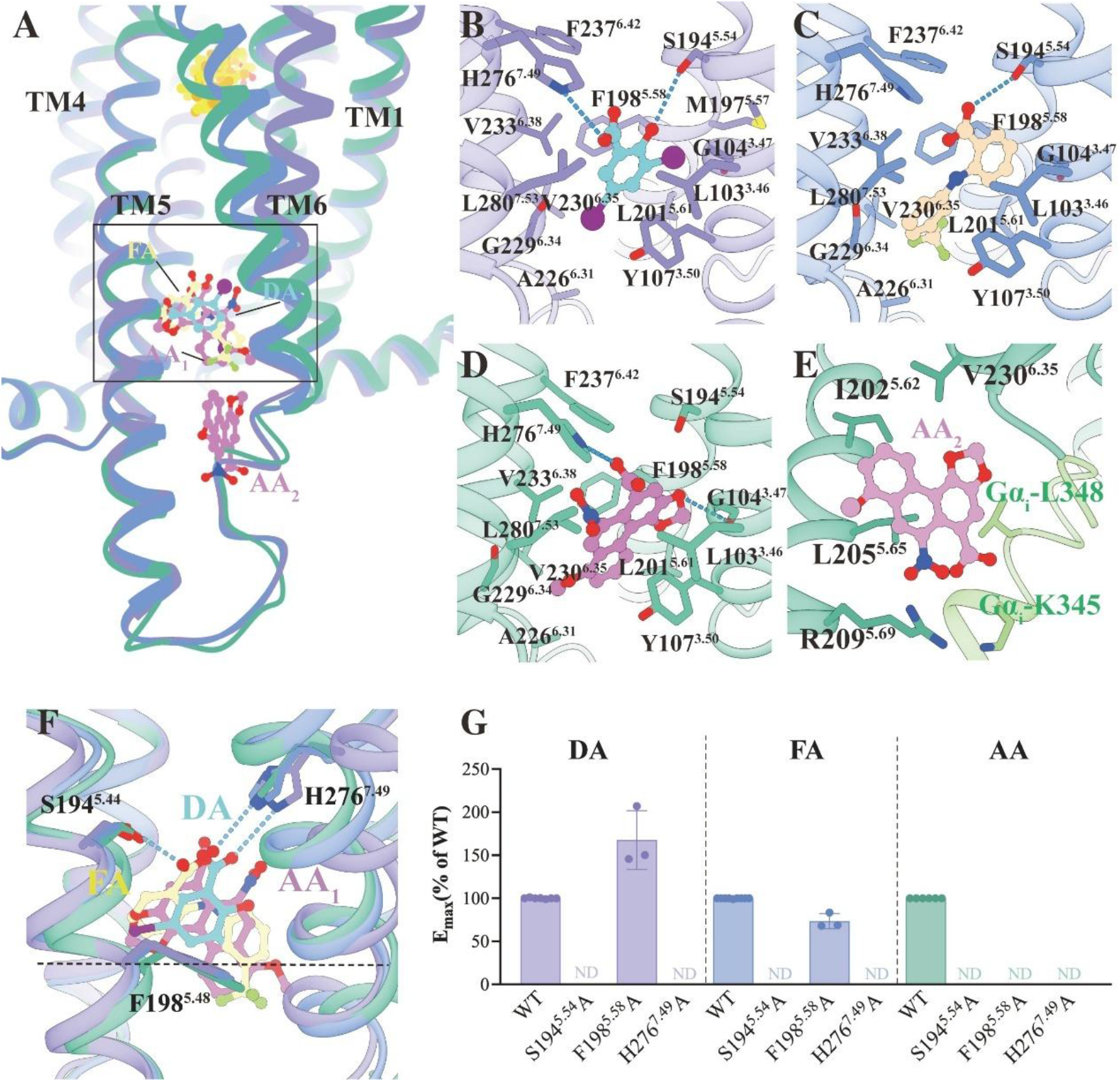
DA/FA/AA binding mode of TAS2R14. (**A**) Comparison of the structures among the DA-bound TAS2R14, FA-bound TAS2R14 and AA-bound TAS2R14. (**B-E**) Amino acids involved in the interactions of DA (B), FA (C), AA_1_ (D) and AA_2_ (E) with TAS2R14. (**F**) Detailed interactions of DA, FA and AA_1_ with TAS2R14. The polar interactions are indicated by blue dashed lines. (**G**) DA-, FA- and AA-induced Ca^2+^assay responses in wild-type (WT) TAS2R14 and mutants in binding pocket. The numbering adopts Ballesteros-Weinstein nomenclature of receptor residues. Data are shown as mean±SEM from at least three independent experiments performed in technical triplicate.

Consistent with the reported TAS2R14–Gi complexes bound to FA and AA (PDB: 8XQS, 8XQO), our structures exhibit their canonical binding pattern, with conserved hydrogen bonds formed with H276^7·49^ and S194^5·54^ and a peripheral hydrophobic environment (Fig. 2C-F, Fig. S5A-C). The same interaction network is observed in the DA-bound complex characterized in the present study (Fig. 2BB,). The more extensive aromatic systems of FA and AA position them closer to TM3 and to the intracellular part of TM6 (Fig. 2F). In addition, the prominent aromatic core of only AA engages in a specific π–π stacking interaction with F198^5.58^, which is absent in DA and FA (Fig. 2F). This ligand-specific structural engagement rationalizes the selective abolition of AA-induced signaling by the F198^5·58^A mutation, with minimal effects on DA or FA-mediated responses (Fig. 2F, Fig. S6, Table S2).

Beyond this conserved binding framework, AA exhibits a unique structural Charateristic. The additional AA_2_ molecule occupies a subpocket enclosed by TM5, TM6 and Gαi-L348, and directly remodels intracellular loop arrangements as well as the G protein coupling interface (Fig. 2A, E). This AA_2_ interaction is structurally flexible : comparison with the TAS2R14–miniGs/gust complex (PDB: 8XQL) reveals a rotated conformation of AA_2_, and it was absent in the earlier AA-bound TAS2R14– Gi structure (PDB: 8XQO) (Fig. S5D). This structural variability, potentially influenced by the higher ligand concentration employed in our sample preparation, highlights the dynamic and context-sensitive nature of AA_2_ binding, suggesting its role may extend beyond simple occupancy to include allosteric modulation of receptor conformation and G protein coupling efficiency.

### Distinct Orthosteric Binding in TAS2R14

Across the three complex structures, we all observed elongated electron density within the orthosteric binding pockets (Fig. S7). Consistent with previous studies, this density aligned well with CHL—a known orthosteric agonist of TAS2R14—in the FA- and AA-bound forms (Fig. 3E-F, Fig. S7B-C). In the DA-bound state, however, this density was larger and penetrated deeper between TM5 and TM6 (Fig. 3C-F, Fig. S7). Considering that a cholesterol analog, cholesterol hemisuccinate (CHS), was supplemented during protein purification, we next performed calcium mobilization assays to explore its potential functional role (Fig. 3A). The results showed that CHS activates TAS2R14 under basal conditions, and elicits a dose-dependent response only after cholesterol depletion with MβCD (Fig. 3B, Fig. S8). The distinct density observed in our structure was unambiguously attributed to CHS. Combined structural and functional evidence collectively demonstrates that CHS acts as an orthosteric agonist, which preferentially stabilizes the receptor in the DA-coupled conformational state, even in the presence of trace CHL contamination (Fig. S8).

**Fig. 3.**
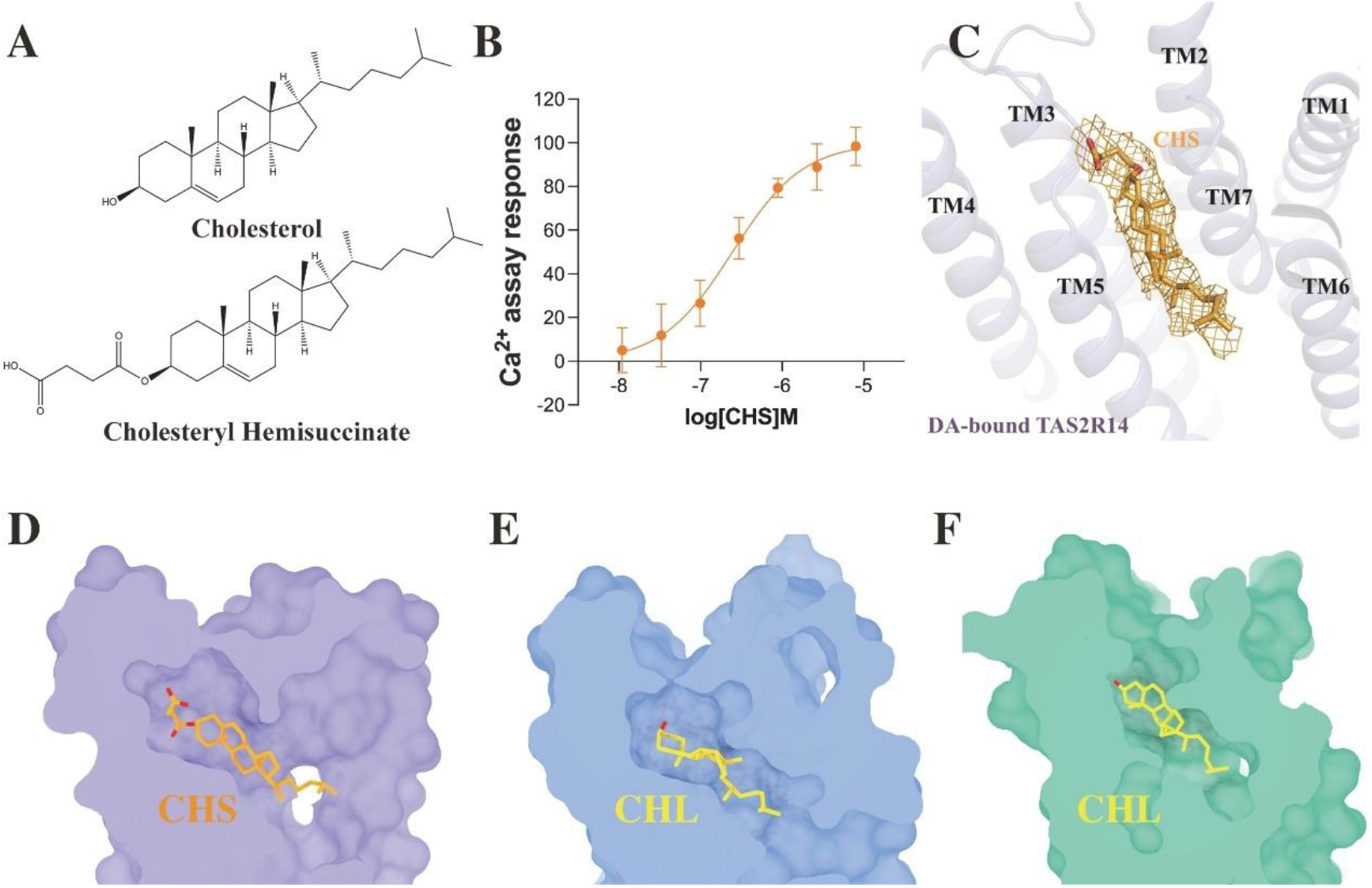
CHL and CHS binding mode of TAS2R14. (**A**) Chemical scaffolds of orthosteric ligands for structural determination. (**B**) The activity of CHS measured in the Ca^2+^mobilization assay. (**C**) The density map of CHS in DA-bound TAS2R14. (**D-F**) Configurations of the orthosteric pocket cross section with CHS in DA-bound TAS2R14(D), with CHL in FA-bound TAS2R14(E), and with CHL in FA-bound TAS2R14(F). Data are mean ± s.e.m. of three independent experiments performed in triplicate.

### Agonist-dependent conformational rearrangements of TM6 in TAS2R14

TAS2R14’s TM6 undergoes significant conformational changes both between the exogenous ligand-free state and agonist-bound states, and among distinct agonist-bound states (Fig. 4A, Fig. S4, Fig. S9).

**Fig. 4.**
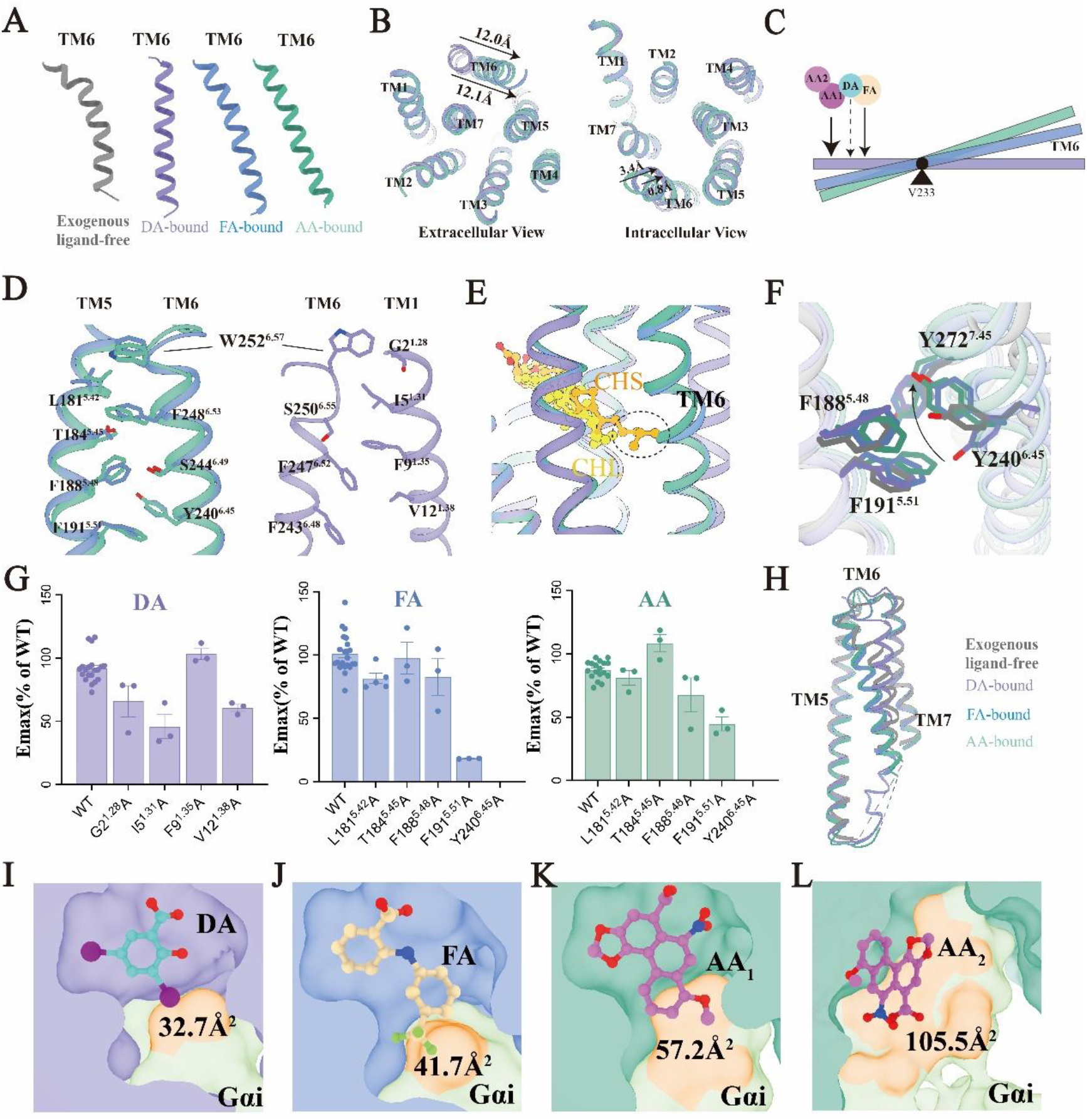
Ligand-specific conformational dynamics in the TAS2R14. (**A**) The conformations of TM6 within exogenous ligand-free TAS2R14 (PDB: 8XQT), DA-bound TAS2R14, FA-bound TAS2R14, and AA-bound TAS2R14. The exogenous ligand-free TAS2R14 (PDB: 8XQT) is shown in gray, DA-bound TAS2R14 in violet, FA-bound TAS2R14 in royal blue, AA-bound TAS2R14 in green. (**B**) Extracellular and intracellular views of DA/FA/AA-bound TAS2R14 structural comparison. (**C**) Schematic diagram of the “lever-like” movement of TM6. (**D**) Hydrophobic network between TM5 and TM6 in FA/AA bound TAS2R14 structures (left), hydrophobic network between TM1 and TM6 in DA-bound TAS2R14 structure (right). (**E**) Comparison of the orthosteric pocket among the DA-bound TAS2R14, FA-bound TAS2R14 and AA-bound TAS2R14. (**F**) Comparison of F188^5.48^, F191^5.51^, Y240^6.45^, and Y272^7.45^ in exogenous ligand-free TAS2R14 (PDB: 8XQT), DA-bound TAS2R14, FA-bound TAS2R14, and AA-bound TAS2R14. (**G**) DA-, FA- and AA-induced Ca^2+^ assay responses in wild-type (WT) TAS2R14 and mutants in interfaces among TM1, TM5, and TM6. (**H**) Comparison of the TM5-7 in exogenous ligand-free TAS2R14 (PDB: 8XQT), DA-bound TAS2R14, FA-bound TAS2R14, and AA-bound TAS2R14. (**I**) Contact area between DA, FA, AA_1_, AA_2_ and G_i1_. Data are shown as mean±SEM from at least three independent experiments performed in technical triplicate.

Interestingly, in the recently reported exogenous ligand-free TAS2R14-Gi complex, TM6 assumes a “curved bow” conformation with its extracellular/intracellular segments proximal to TM5 and mid-region deviating away (Fig. 4A). Along with allosteric agonist DA/FA/AA binding induces a straightening of TM6, which is most pronounced in the DA-bound state (Fig. 4A, Fig. S9). DA’s small structure permits substantial inward displacement of TM6’s cytoplasmic segment; FA’s and AA’s dual or three aromatic rings impose moderate steric constraints, limiting TM6 flexibility (Fig. 4B, Fig. S9); and AA’s dual cytoplasmic binding drives TM6 outward. (Fig. 2A, Fig. 4B). On the cytoplasmic side, the TM6 helix in the DA-bound state is retracted by 0.8 Å and 3.4 Å relative to the FA- and AA-bound states, respectively (Fig. 4B).

Furthermore, this lever-like movement of TM6 directly couples intracellular conformational rearrangements to extracellular changes (Fig. 4C). Notably, structural comparison reveals two distinct extracellular poses of TM6, involving a lever-like movement that displaces the Cα atom of W252^6.57^ by up to 12 Å between the DA-bound and FA/AA-bound states (Fig. 4B). In the FA/AA-bound states, TM6 engages TM5, forming a network of hydrophobic contacts (Fig. 4D, F-G, Fig. S10), which imposes steric constraints on CHS, precluding its stable occupancy of the orthosteric pocket (Fig. 4A-E).

In contrast, in the DA-bound state, the extracellular end of TM6 swings toward TM1, and W252^6.57^ flips 180º to form a unique TM1-TM6 interface (Fig. 4D, F-G, Fig. S10), losing the contact with TM5. This conformation induces a unique cleft between TM5 and TM6 that has not been identified in previous TAS2R family structures, and this distinctive cavity accommodates CHS with adequate spatial compatibility (Fig. 4E, 4H).

### Molecular Mechanism underlying the differential ligand efficacies of TAS2R14

An efficacy ranking (E_max_) of AA > FA > DA on TAS2R14 originates from their inherent structural divergences, which sequentially mediate TM6 conformational changes, reshape the cytoplasmic cavity, and ultimately modulate G protein coupling efficiency (Fig. 1B, Table S4).

There are two key factors. First, DA/FA/AA binding remodels the TM6 conformation, leading to the selective stabilization of distinct TM interfaces (Fig. 4A, D). In the FA-and AA-bound state, there is an aromatic cage composed of F188^5·48^, F191^5·51^, Y240^6·45^, and Y271^7·45^ that stabilizes middle TM5-TM6-TM7 interface, whereas in DA-bound state, the Y240^6·45^ swing down and aromatic cage is absent (Fig. 4F), and the extracellular part of TM6 form new contact with TM1 (Fig. 4D). Interestingly, the aromatic cage is also absent in ligand-free with low activity. In addition, Mutagenesis of Y240^6·45^A that disrupts TM5-TM6-TM7 interface abolished agonist-induced signaling, while disrupting TM1-TM6 had minimal effect on DA (Fig. 4G, Fig. S10, Table S2). That indicates the middle TM5-TM6-TM7 interface is critical for receptor activation. Second, DA, FA and AA all participate directly in the formation of the cytoplasmic cavity, with interaction surfaces of 32.9 Å^2^, 41.7 Å^2^and 54.8 Å^2^+105.5 Å^2^with the α5 helix of G protein, respectively (Fig. 4I-L). Larger interaction interfaces correlate with stronger G protein engagement, contributing to higher efficacy. This combined regulation of the receptor-G protein interface directly tunes downstream signaling output, ultimately explaining the graded efficacies of DA, FA, and AA.

### Orthosteric–Allosteric Coupling Modulation

The identification of CHL and CHS as orthosteric agonists and of DA, FA, and AA as allosteric agonists for TAS2R14, combined with functional signaling assays, reveals a intricate, diversified orthosteric–allosteric coupling mechanism that differentially modulates agonist potency.

This complexity is evident in three key observations. First, the depletion of CHL using MβCD revealed an orthosteric-to-allosteric coupling effect between CHL and the three allosteric agonists, DA, FA and AA. As shown in Figure 4 A-F, this depletion resulted in a reduction in the potencies of DA, FA, and AA by approximately 1.2-fold, 1.8-fold, and 3.6-fold, respectively (Fig. 5A-F, Table S3). Second, the addition of CHS demonstrated an orthosteric-to-allosteric coupling with DA, where the potency of DA was significantly amplified by about 5-fold with CHS addition in the context of CHL depletion, and by 2.3-fold when CHS was added without prior CHL depletion (Fig. 5A-F, Table S3). Third, in contrast, the addition of CHS had no effect on the potencies of the other two allosteric agonists, FA and AA, suggesting an orthosteric-to-allosteric uncoupling between CHS and these agonists (Fig. 5A-F, Table S3).

**Fig. 5.**
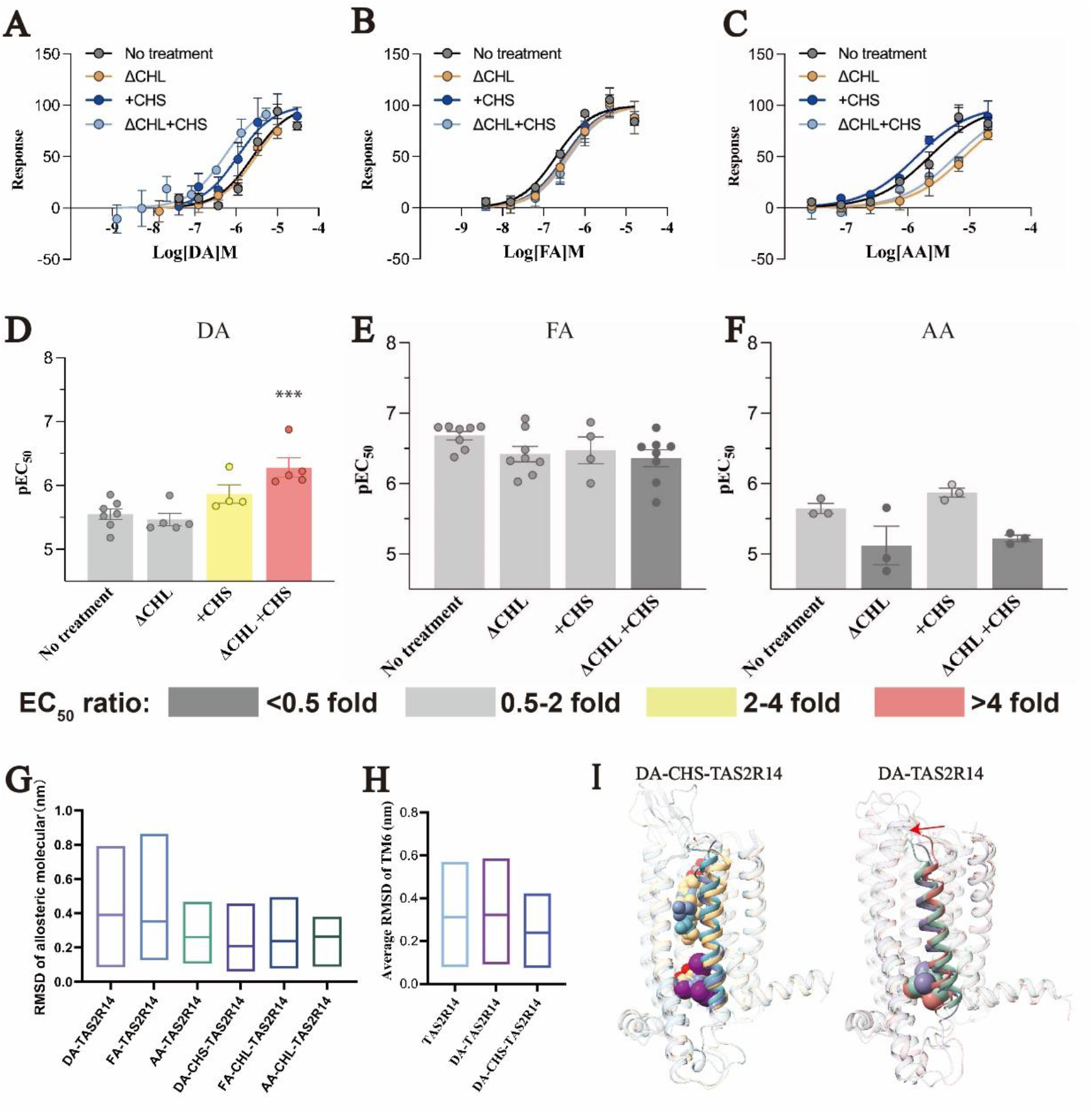
Orthosteric-to-allosteric coupling modulate mechanism. (**A-C**) Ca^2+^mobilization assay responses in no treatment TAS2R14 stimulated by DA, FA and AA in the presence of CHS or CHL relative to reference the three allosteric agonists alone. (**D-F**) pEC_50_ of three allosteric agonists with or without CHS measured by signaling assay. (**G-H**) The average r.m.s.d. values of allosteric molecules (G) and TM6 (H) during molecular dynamic simulation. (**I**) The superposition of the representative frames extracted from DA-CHS-TAS2R14 system (left) and DA- TAS2R14 system (right). DA and CHS are shown as sphere. Data are shown as mean±SEM from at least three independent experiments performed in technical triplicate.

Molecular dynamics (MD) simulations further clarify the delicate the fine-tuned regulation of orthosteric-to-allosteric coupling. Both orthosteric ligands, CHL and CHS, restrict the structural dynamics of TAS2R14 in complex with DA, FA or AA (Fig. 5G, Fig. S11-12). Notably, DA exhibits greater conformational flexibility than FA and AA in the absence of orthosteric ligands (Fig. 5G, Fig. S11-12). Furthermore, simulations of ligand-free TAS2R14 and DA-bound TAS2R14 revealed a significant elevated TM6 r.m.s.d. values, whereas simultaneous binding of DA and CHS effectively limited TM6 fluctuation (Fig. 5H, Fig. S11-12). Moreover, the extracellular segment of TM6 tends to approach TM5 in the absence of orthosteric ligand CHS (Fig. 5I). These MD simulation results suggest that CHS and DA synergistically stabilize the TM6 conformation, thus regulating TAS2R14 activity. These findings are consistent with our biochemical experiment data, which demonstrate a significant coupling mechanism between DA and CHS as distinct from the behavior observed with FA and AA.

## Discussion

The mammalian bitter taste receptor TAS2R14 represents a remarkable paradigm of G protein-coupled receptor (GPCR) functional and structural plasticity, uniquely capable of recognizing an extraordinarily diverse chemical repertoire of xenobiotic toxins, natural bitter compounds, and clinical pharmaceuticals. Decades of sensory biology and GPCR pharmacology research have long puzzled over the fundamental mechanistic question of how a single TAS2R subtype achieves such exceptional ligand promiscuity while translating structurally distinct agonist inputs into strictly graded, ligand-specific signaling efficacies. Prior cryo-electron microscopy (cryo-EM) studies of TAS2R14 have identified rudimentary orthosteric cholesterol-binding pockets and intracellular allosteric ligand-binding sites, yet the critical conformational interplay linking differential agonist engagement, transmembrane helix rearrangement, G protein coupling efficiency, and cross-talk between discrete receptor binding cavities has remained poorly resolved and mechanistically uncharacterized. Here, our integrative high-resolution cryo-EM structural determination, functional calcium mobilization assays, site-directed mutagenesis validation, and MD simulation analyses collectively delineate a sophisticated orthosteric–allosteric dual-coupling regulatory mechanism governing TAS2R14 ligand recognition, receptor activation, and graded signaling output, filling a longstanding critical gap in the structural and mechanistic understanding of bitter taste receptor promiscuity and GPCR ligand-directed functional bias.

We propose that the receptor exploits malleable orthosteric and allosteric pocket geometries, coupled with pronounced TM6 conformational flexibility, to expand ligand compatibility. Differential TM6 remodeling at intracellular and extracellular regions dynamically reshapes both binding cavities, allowing the receptor to tolerate sterols with variable carbon-chain lengths, including the longer-chain CHS (Fig. S13). This interplay between pocket plasticity and TM6 dynamics therefore provides a plausible mechanistic explanation for the exceptional ligand promiscuity of TAS2R14.

We address the unresolved molecular question by elucidating the precise structure–activity relationship (SAR) that underpins graded agonist efficacy at TAS2R14, with the conformational plasticity of TM6 serving as the central regulatory hub that translates intrinsic chemical properties of distinct bitter ligands into quantitatively differentiated receptor signaling outputs. High-efficacy AA and FA specifically stabilize a conserved aromatic network centered on the key residue Y240^6.45^ at the TM5-TM6-TM7 bundle to sustain full receptor activation; by contrast, both low-efficacy DA and the ligand-free baseline receptor fail to maintain this Y240^6.45^-dependent activating aromatic cage, which disrupt hydrophobic packing to restrict activation. Complementing this helical rearrangement, agonists also differ substantially in their contact interfaces with the G protein α 5 helix, with larger interaction surfaces corresponding to stronger G protein engagement and higher functional efficacy.

Together, these results establish a straightforward mechanistic model explaining how structural variations among agonists dictate TAS2R14 signaling efficacy.

Following the structural principles governing ligand recognition and graded efficacy, our work reveals that TM6 conformational plasticity underpins the selective coupling crosstalk between orthosteric sterols and allosteric agonists. Agonist-specific TM6 structural states dictate the permissiveness of orthosteric sterol binding, thereby determining whether allosteric modulation can occur. This conformational selection explains the preferential potentiation of DA, but not FA and AA, by CHS, highlighting a ligand-specific coupling mode intrinsic to TAS2R14. This mechanism accounts for the specific potentiation of DA potency by CHS without affecting FA and AA, demonstrating the inherent ligand- and sterol-dependent coupling selectivity of TAS2R14. Such regulatory asymmetry is physiologically meaningful, as sterol-dependent orthosteric tuning enables dynamic adjustment of receptor sensitivity to varying bitter stimuli. The orthosteric–allosteric tuning allows TAS2R14 to act as a finely controlled sensory switch that dynamically adjusts receptor sensitivity according to endogenous sterol levels and local ligand microenvironments, thereby shaping bitter taste sensitivity, maintaining metabolic homeostasis, and potentially influencing inflammatory and pathological processes.

In summary, the present study resolves key longstanding mechanistic uncertainties regarding TAS2R ligand promiscuity, graded efficacy, and selective orthosteric – allosteric modulation, with TM6 conformational dynamics identified as the unifying structural determinant governing all core receptor regulatory properties. These findings refine the current understanding of bitter taste receptor activation and sterol-dependent tuning, providing a fundamental structural and functional framework for future exploration of TAS2R-related physiological functions and targeted modulator development.

## Materials and Methods

### Construct cloning

To obtain the agonist bound TAS2R14-G_i_ complex, the human TAS2R14 gene was cloned into a modified pFastBac1 vector with the haemagglutinin signal peptide (HA) signal peptide at N terminus, and a Flag tag (DYKDDDD) and a PreScission protease site, and a twin-Strep-tag (WSHPQFEK-GGGSGGGSGGSA-WSHPQFEK) at C terminus. To enhance protein yield and stability, bRIL was fused at the N terminus and 18 residues at C terminus of TAS2R14 were truncated. Moreover, two mutations M205^5.65^L and G283^7.56^T were introduced into the construction, using standard QuikChange polymerase chain reaction (PCR). The dominant-negative human Gα_i1_ (DNGα_i1_) was generated as previously described ^23^ by introducing four mutations, S47N, G203A, A326S, and E245A. To obtain apo-TAS2R14-G_i_ complex, the NanoBiT strategy was employed with an additional mutation S25^1.51^G to enhance the stability of the complex ^24^. DNGα_i1_, human Gβ1, Gγ2, and a single chain antibody scFV16 ^25^ were cloned into the pFastBac vector.

### Insect cell expression

Coexpression of receptor TAS2R14, DNGα_i1_ and Gβ1γ2 were conducted in HighFive insect cells (Invitrogen) using the Bac-to-Bac Baculovirus Expression System (Invitrogen). Cells were grown to a density of 3×10^6^ cells per milliliter and then coinfected with the high-titer recombinant baculovirus at a multiplicity of infection (MOI) ratio of 1:1:1. The cells were collected by centrifugation after transfection for 48 hours at 27°C, and stored at -80 °C until use.

### Purification of apo/DA/FA/AA-TAS2R14-G_i_ complexes

Cells were thawed and suspended in a buffer containing 20 mM HEPES (pH 7.5), 50 mM NaCl, 2 mM MgCl_2_ and EDTA-free protease inhibitor cocktail tablets (Roche). The formation of the TAS2R14-Gα_i_ complex was facilitated by adding 10 μg/ml ScFv16 prepared as previously reported ^26^, and 25 mU/ml Apyrase (New England BioLabs) to the cell suspension. Simultaneously, individual additions of 400 μM 3,5-Diiodosalicylic acid (DA, Sigma, D124001), 250 μM Flufenamic acid (FA, Sigma, F9005), and 400 μM Aristolochic acid A (AA, Bidepharm, BD16542) were made to obtain agonist-bound TAS2R14-Gαi complexes. These suspensions were then incubated at 16 °C for 1 hour. The membrane pellets were collected by ultracentrifugation at 38,000 rpm for 30 minutes. The complexes were extracted from the membrane with an extraction buffer containing 20 mM HEPES (pH 7.5), 150 mM NaCl, 2 mM MgCl_2_, 0.5% (w/v) lauryl maltose neopentyl glycol (L-MNG, Anatrace), 0.025% (w/v) cholesteryl hemisuccinate (CHS, Anatrace), 25 mU/ml Apyrase, and 400 μM DA/250 μM FA/400 μM AA. After a 2-hour incubation period at 4 °C, the supernatant was isolated via ultracentrifugation at 38,000 rpm for 30 minutes, and subsequently incubated overnight with Strep-Tactin®XT 4Flow® resin (IBA) at 4 °C. The resin was then washed with 20 column volumes of wash buffer containing 25 mM HEPES (pH 7.5), 150 mM NaCl, 2 mM MgCl_2_, 0.01% (w/v) L-MNG, 0.0005% (w/v) CHS, and 400 μM DA/250 μM FA/400 μM AA. After which, the resin was eluted with 10 column volumes of 25 mM Tris (pH 7.5), 150 mM NaCl, 2 mM MgCl_2_, 50 mM Biotin, 0.01% (w/v) L-MNG, 0.0005% (w/v) CHS and 400 μM DA/250 μM FA/400 μM AA. The complexes were further purified by size-exclusion chromatography using a Superdex 200 Increase 10/300 column (GE Healthcare) pre-equilibrated with 20 mM HEPES (pH 7.5), 150 mM NaCl, 2 mM MgCl_2_, 0.01% (w/v) L-MNG, 0.0005% (w/v) CHS and 400 μM DA/250 μM FA/400 μM AA. The same protocol was used for purification the apo form, excluding the addition of an agonist. Finally, the purified complexes were concentrated to 3-4 mg/ml with a 100-kDa molecular weight cut-off concentrator (Millipore), and were analyzed by SDS-PAGE.

### Cryo-EM grid preparation and data collection

For the preparation of cryo-EM grids, 3μl of the purified complex samples were applied onto freshly glow-discharged holey carbon grids (Quantifoil R1.2/1.3, Au 300mesh). These grids were subsequently blotted for 3 seconds in conditions of 100% humidity at 4°C before being rapidly plunge-frozen in liquid ethane using Vitrobot Mark IV (FEI). The cryo-EM images were recorded through the EPU software on a 300kV Titan Krios G3i electron microscope (FEI) with a K3 summit direct electron detector and a Gatan Quantum energy filter at a magnification of 105,000×, corresponding to a pixel size of 0.851Å. Each micrograph was then dose-fractionated with 40 frames for 2.5 seconds to accumulate to a total dose of 54 e^-^Å^-2^, with a defocus range of -1.0 to -1.5μm.

### Cryo-EM data processing

For the DA- and FA-TAS2R14-G_i_ complex, a total of 5,107 and 3,858 movies respectively underwent beam-induced motion correction and dose-weighting using RELION MotionCor2 ^27^. Contrast transfer function estimation was performed using Gctf ^28^. Cryo-EM data processing was subsequently carried out with CryoSPARC ^29^. For initial reference, 800 movies were randomly selected for each batch, which were then followed by blob picker, ab-initio reconstruction and heterogeneous refinement. The good classes were processed with non-uniform refinement and used to create templates for further template picker. A total of 4,987,646 and 3,812,717 particles were extracted respectively and subjected to several rounds of ab-initio reconstruction and heterogeneous refinement. The best classes underwent 2D class processing. A subset of 1,385,801 and 1,296,323 particles were chosen for further non-uniform refinement followed by local resolution estimation with the final resolution of DA- and FA-TAS2R14-G_i_ complex at 2.44 Å and 2.39 Å, respectively. Signal subtract was performed for better density of the trans-membrane domain of DA- and FA-TAS2R14 using RELION and a final resolution of 3.13 Å and 2.98 Å was generated after a series process of Refine3D and postprocess.

For AA-TAS2R14-G_i_ complex, 4,429 movies were recorded for data processing, respectively. A total of 4,270,697 particles were extracted using the density map of DA-TAS2R14-G_i_ complex as initial reference. Several rounds of ab-initio reconstruction and heterogenous refinement were performed. For AA-TAS2R14-G_i_ complex, the well-defined class with 1,474,057 were selected for further 2D class and a subset of 1,166,258 particles was then subjected for following non-uniform refinement and local resolution estimate with a final resolution of 2.69 Å at a Fourier shell correlation of 0.143. A resolution of 3.17 Å of the trans-membrane domain of AA-TAS2R14-G_i_ complex was obtained after signal subtract, refine3D and postprocess.

For apo-TAS2R14-G_i_ complex, 4,064 micrographs were collected and 800 micrographs were randomly selected for initial reference. A total of 4,062,875 particles were extracted for two rounds of ab-initio reconstruction and heterogenous refinement. The well-defined class with 937,135 particles were subjected to 2D class and 762,384 particles were selected for non-uniform refinement and local resolution estimation with the final resolution of 2.68 Å.

### Model building and refinement

For the structure of the DA-TAS2R14-G_i_ complex, the initial template of TAS2R14 was generated using AlphaFold2^30^. DA and CHS coordinates and geometry restrain were generated using Phenix^31^, elbow. Models was docked into the density map using UCSF Chimera^32^, followed by iterative manual adjustments in COOT^33^. The model was further refined by real-space refinement using Phenix.

For FA-, AA- and apo-TAS2R14-G_i_ complex, the coordinates of TAS2R14 and G_i_ from DA-TAS2R14-G_i_ complex were used as initial model. FA,AA and CHL coordinates and geometry restrain were generated using phenix, elbow. Each model was rigidly docked into density map in Chimera and then manual adjusted using COOT, followed by iterative refinement using Phenix. The final refinement statistics for all the structures were performed using Phenix and figures of the structures were generated using UCSF Chimera.

### Intracellular calcium mobilization assay

We constructed stable transfected cell lines expressing Gα_16/gust44_ ^34^ using the Flp-In™ T-REx™ 293 cell line (Thermo Fisher, R78007) with its corresponding plasmids pcDNA™5/FRT/TO (Thermo Fisher, V652020) and pOG44 (Thermo Fisher, V600520), following the instructions provided in the manual. The G_α16/gust44_cell lines were cultured in complete DMEM containing 10% (v/v) fetal bovine serum ((FBS Gibco, 10270106)) and 1% penicillin-streptomycin (Gibco, 15140122). The cells were seeded at a density of 3×10^6^ cells per well in 6 cm culture dishes and incubated at 37°C overnight. Then the cells were transiently transfected with 4 μg Wild-type TAS2R14 or mutant TAS2R14 plasmid DNA using Polyethylenimine Linear (PEI) MW40000 (YEASEN, 40816ES03). After 6-8 hours of transfection, the medium was replaced with complete DMEM containing 2 μg/ml puromycin. 24 hours after transfection, the cells were diluted in complete DMEM containing 2 μg/ml doxycycline (Solarbio, D8960) and seeded at a density of 80,000 cells per well in a pre-coated black clear-bottom transparent 96-well plate (Corning®, 3340) with D-Lysine homopolymer hydrobromide (Sigma, P7886). 24 hours later, the medium was replaced by 40 μL HBSS buffer containing 20 mM HEPES (pH 7.4), 4 mM Fluo-8H™, and 2.5 mM probenecid. The plate was then incubated at 37°C for 40 minutes. Finally, the baseline was measured and using a FlexStation 3 Multi-Mode Microplate Reader at 37°C in 100 μL HBSS buffer containing 20 mM HEPES (pH 7.4). At 20 seconds, the instrument was set to automatically add 50μL of a triple concentration of DA/FA/AA/3W, and the reading time continued for 120 seconds (data was collected every two seconds). The data were analyzed using GraphPad Prism 9.0 through nonlinear regression analysis. Wild-type TAS2R14 and mutant TAS2R14 were cloned into the pcDNA3.1 vector and tagged with HA and Flag tags at the N-terminus of the receptors (as mentioned above). Empty pcDNA3.1 vector was used as a negative control. The cell surface expression level was detected by monoclonal anti-FLAG M2-fluorescein isothio-cyanate antibody (Sigma-Aldrich).

In the CHS activation of T2R14 and synergistic experiments, cells transfected for 24 hours were transferred to a 96-well plate and cultured in DMEM supplemented with 1% FBS, 1% penicillin-streptomycin, and 2 μg/ml doxycycline. Due to the low solubility of CHS, it was dissolved in HBSS containing 5 mM Methyl-β-cyclodextrin (MβCD; Sigma, C4555). For experiments necessitating cholesterol depletion, the cells were incubated in HBSS with 5 mM MβCD at 37°C for 1 hour before incubating the dye. In the synergistic experiments, to ensure simultaneous addition of DA/FA/AA and CHS to the detection plate, various concentrations of DA/FA/AA were mixed with CHS, maintaining a final concentration of CHS at 5 μM. The remaining experimental protocols were consistent with those previously outlined.

### Molecular dynamics simulation

First, system construction was initiated. All systems were constructed using the membrane builder module of charmm-gui ^35^. The systems were contained within cubic boxes of dimensions 80Å×80Å×150Å. Each system comprised a bilayer membrane of 1-palmitoyl-2-oleoyl-sn-glycero-3-phosphocholine (POPC), modeled with TIP3P ^36^ water molecules at a concentration of 0.15 M NaCl salt ions, along with protein and small molecule complexes. The charmm36 ^37^ force field was applied uniformly across all systems.

Gromacs2021 ^38^was employed for simulations of all systems. Each system underwent energy minimization and pre-equilibration. Energy minimization was conducted using steepest descent with gradual relaxation. Pre-equilibration employed the isothermal-isobaric (NPT) ensemble, with temperature control via the v-rescale ^39^ method and pressure control using the Berendsen barostat ^40^.

Eventually, production simulations of 300 ns were conducted in every systems. The systems were maintained at a stable temperature of 310 K, controlled by Parrinello-Rahman coupling to 1 atmosphere, with a coupling parameter τp set to 1.0 ps. SETTLE ^41^ and LINCS ^42^ methods were applied to constrain water molecules and other molecules involving hydrogen atoms in covalent bonds, respectively. A simulation time step of 2 fs was utilized, and electrostatic interactions were computed using the Particle-Mesh Ewald (PME) ^43^ algorithm with a cutoff distance of 1.4 nm.

## Funding

This work was supported by in part by Ministry of Science of Technology (2023YFA0916300 to A.Q. and 2020YFA0908500 to S.Y.), the National Natural Science Foundation of China (32271249 to A.Q. and 31971127 to S.Y.), The Distinguished Young Scholars of Hubei Province (2022CFA078 to S.W.), and Knowledge Innovation program of Wuhan-Shugung Project (2023020201020418 to S.W.).

## Author contributions

T.Z. optimized the constructs, developed the expression and purification procedures, prepared the protein samples for cryo-EM, and performed SPR and signaling assay; J.W performed negative-stain EM data acquisition and analysis, cryo-EM data processing and analysis, model building, and structure refinement; Z.O. helped with cryo-sample preparation; Z.O. and Y.Z. helped perform signaling assay and data analysis; S.W. oversaw EM data acquisition, analysis, processing and helped with data analysis and edited the manuscript; A.Q. and S.Y. initiated the project, planned and analyzed experiments, supervised the research, and wrote the manuscript with input from all co-authors.

## Competing interests

The authors declare no competing interests.

## Data and materials availability

Atomic coordinates and the cryo-EM density maps for structures of DA-TAS2R14-Gi1, FA-TAS2R14-Gi1, AA-TAS2R14-Gi1, FA-TAS2R14-miniGgust, and apo-TAS2R14-Gi1 have been deposited in the RCSB Protein Data Bank (PDB) with identification codes XXXX, XXXX, XXXX, XXXX and XXXX, and EMD-XXXX, EMD-XXXX, EMD-XXXX, EMD-XXXX, and EMD-XXXX. All other data are available in the manuscript or the supplementary materials. Reagents are available from the corresponding authors upon reasonable request.

## Notes

### Competing Interest Statement

The authors have declared no competing interest.

