## Supplementary material for "Structural Basis of Ligand-Specific Orthosteric-Allosteric Coupling and Sensory Tuning in the Human Bitter Taste Receptor TAS2R14": Fig. S & Table S

**Supplementary Materials**

Acknowledgments

We thank Zhiguang Yuchi for providing Flp-In™ T-REx™ 293 cell line, plasmids pcDNA™5/FRT/TO and pOG44.

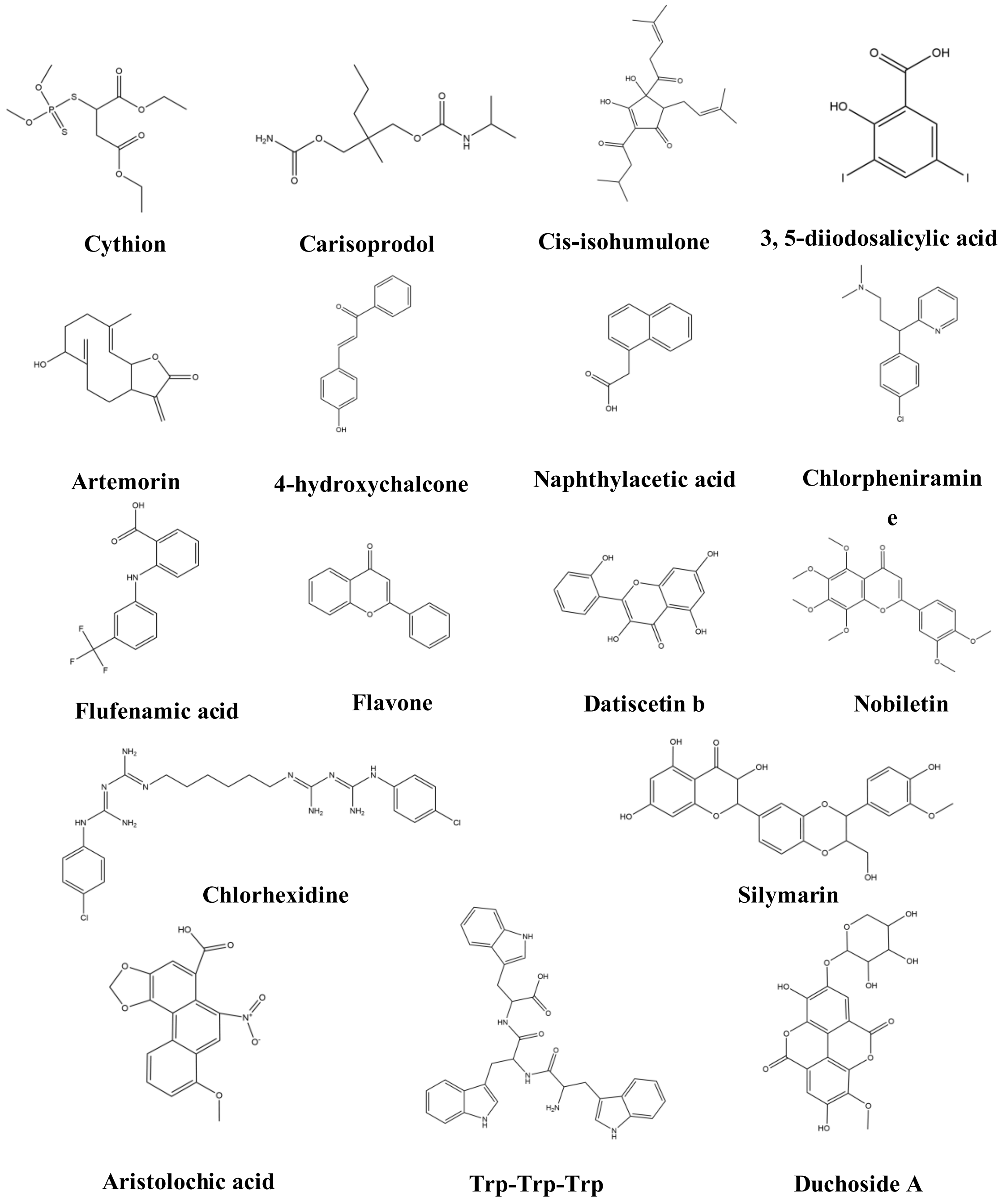


**Fig. S1. Distinct chemical scaffolds of TAS2R14 ligands.**

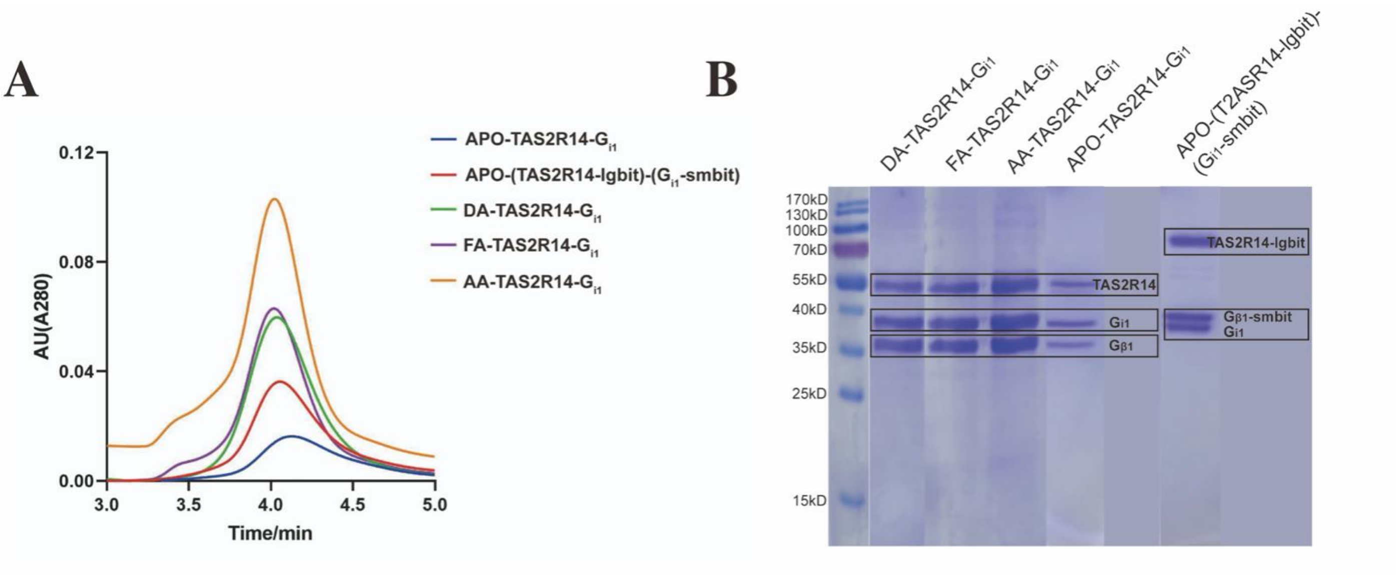


**Fig. S2. Protein quality and extracellular view of apo-TAS2R14-$G_i$ complex.** (**A** to **B**) SEC (A) and gel (B) results of comparison of apo-TAS2R14-$G_i$ with DA/FA/AA-TAS2R14-$G_i$ samples.

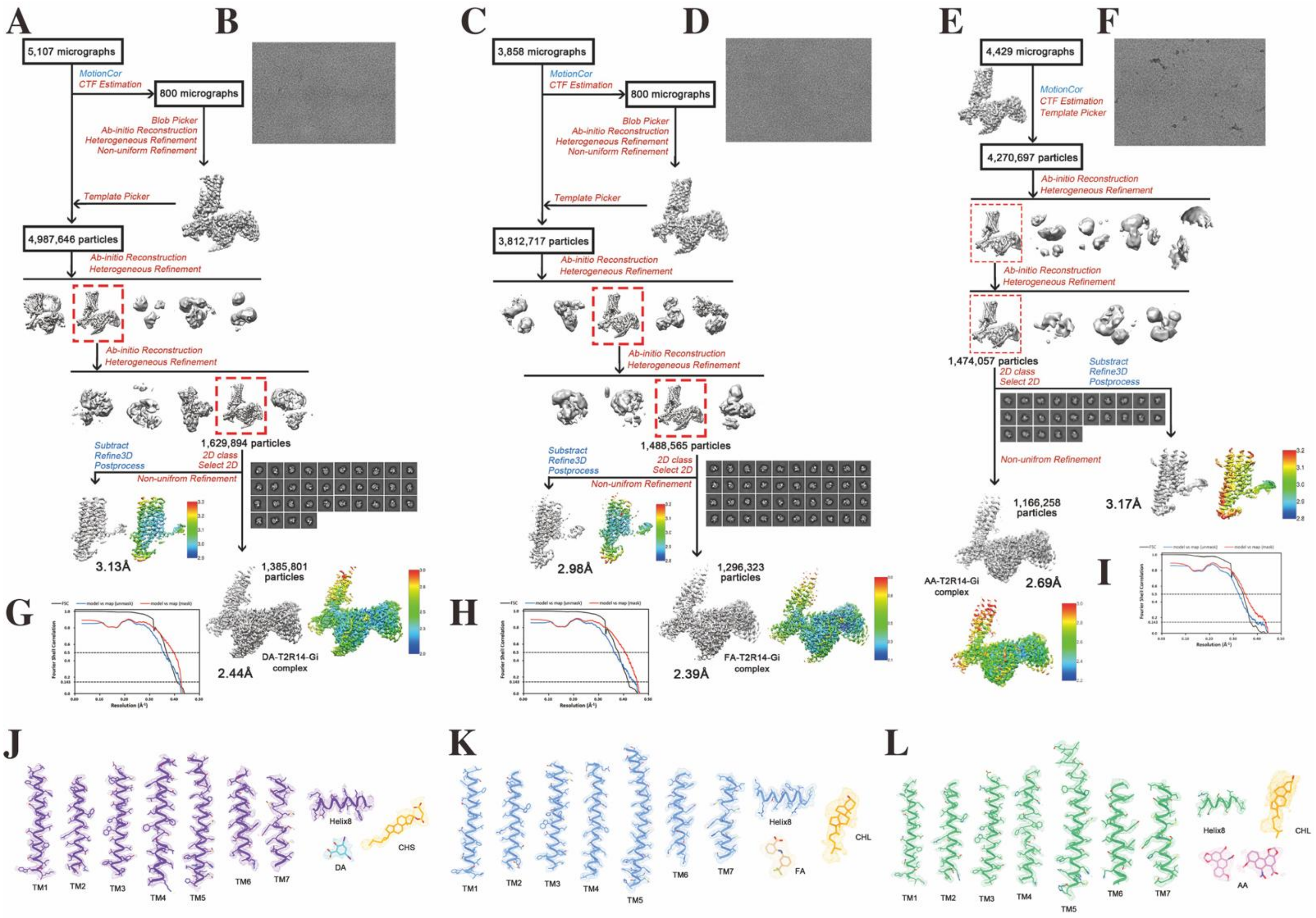


**Fig. S3. Cryo-EM data imaging and processing of the DA-, FA- and AA-TAS2R14-G$_i$ complexes.** (A, B, G, J) Results for the DA-TAS2R14-G$_i$ complex. (C, D, H, K) Results for the FA-TAS2R14-G$_i$ complex. (E, F, I, L) Results for the AA-TAS2R14-G$_i$ complex. (A, C, E) Cryo-EM data process workflow. (B, D, F) Representative micrographs. (G, H, I) Fourier shell correlation (FSC) curves of the density maps. (J, K, L) Cryo-EM density of transmembrane helices and ligands.

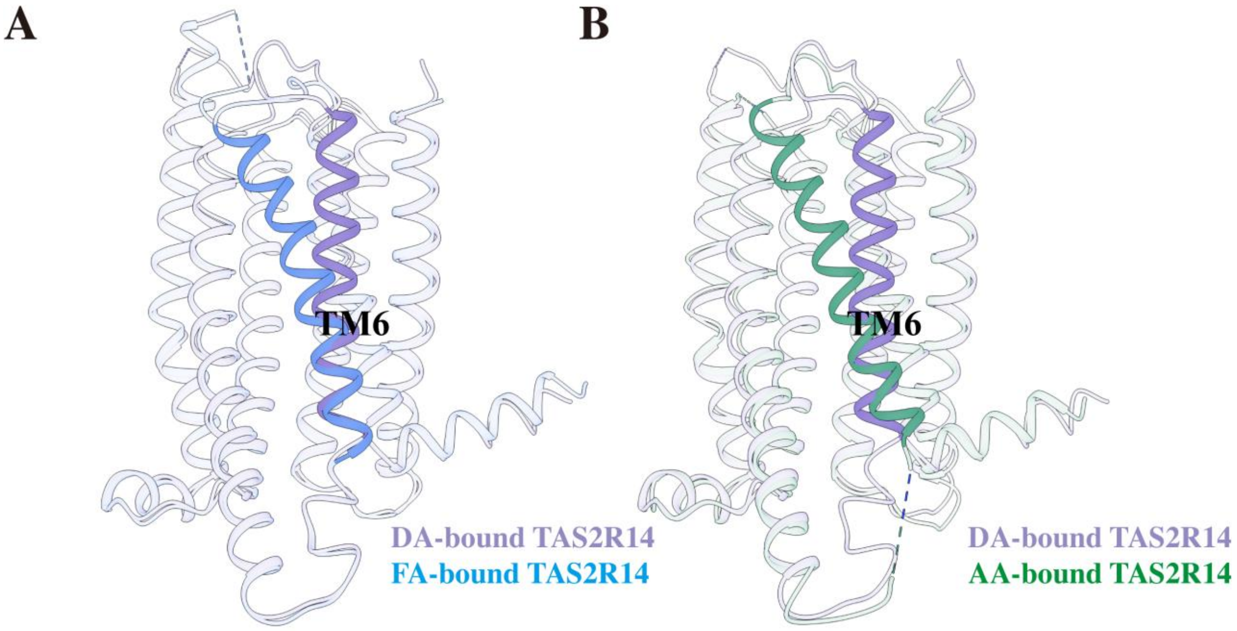


**Fig. S4. Comparisons of the structures of the DA-bound TAS2R14 with FA-bound TAS2R14 (A), and with AA-bound TAS2R14 (B).**

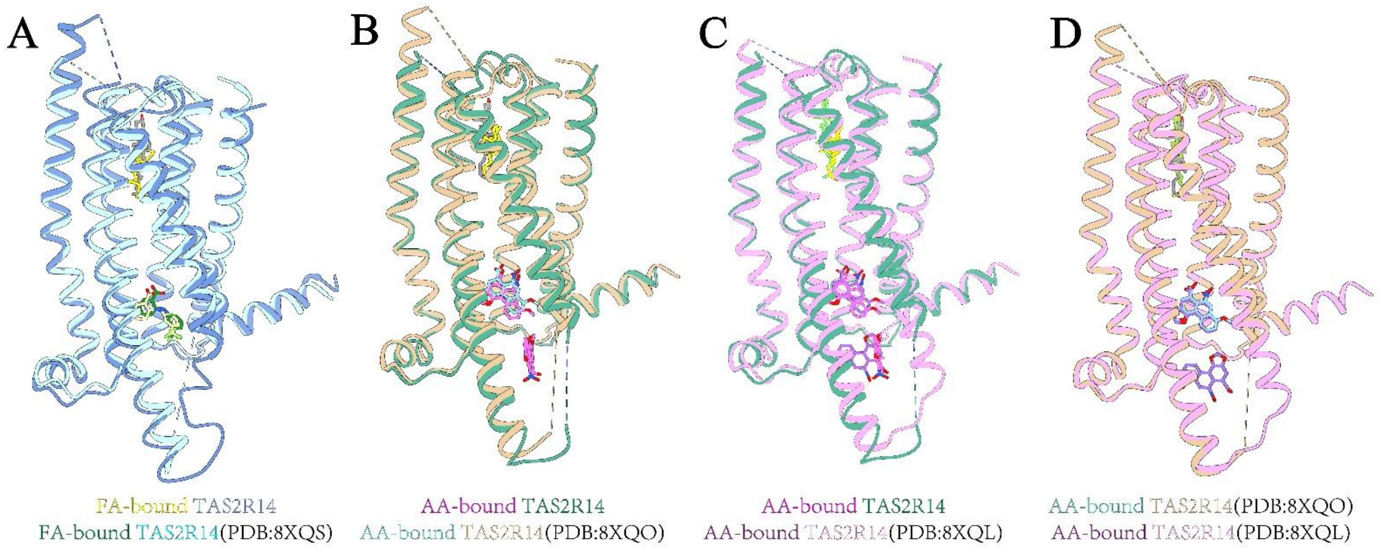


**Fig. S5. Comparisons of different structures of TAS2R14.** (**A**) Comparison of FA-TAS2R14-$G_i$ with FA-TAS2R14-DN$G_i$ (8XQS). The FA-TAS2R14-$G_i$ is shown in royal blue, FA-TAS2R14-DN$G_i$ (8XQS) in cyan. (**B**) Comparison of AA-TAS2R14-$G_i$ with AA-TAS2R14-$G_i$ (8XQO). The AA-TAS2R14-$G_i$ is shown in green, AA-TAS2R14-DN$G_i$ (8XQO) in yellowish. (**C**) Comparison of AA-TAS2R14-$G_i$ with AA-TAS2R14-$G_i$ (8XQL). The AA-TAS2R14-mini$G_{s/gust}$ (8XQL) is shown in light pink. (**D**) Comparison of AA-TAS2R14-mini$G_{s/gust}$ (8XQL) with AA-TAS2R14-$G_i$ (8XQO).

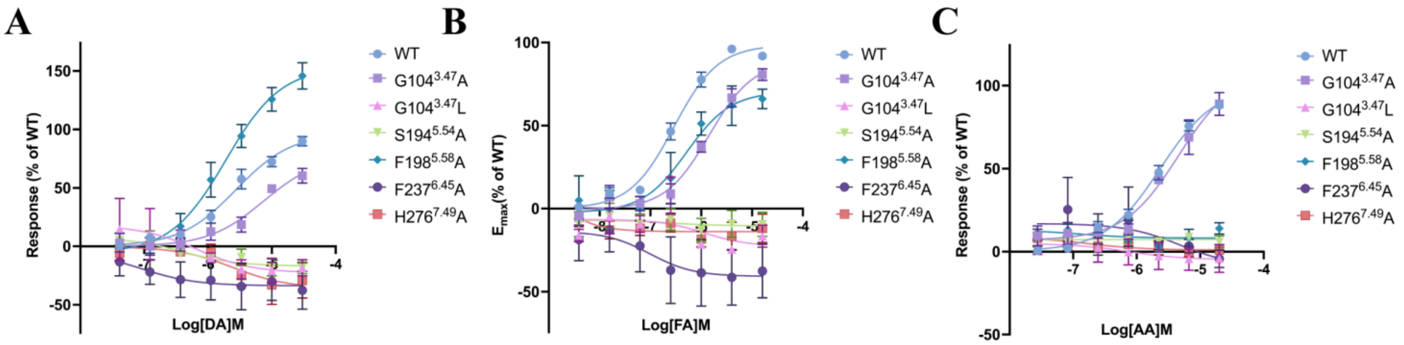


**Fig. S6. Dose responses curves of residue mutations in ligand binding pockets and receptor activation for DA (A), FA (B) and AA (C) in calcium mobilization assay.** Data are shown as mean±SEM from at least three independent experiments performed in technical triplicate.

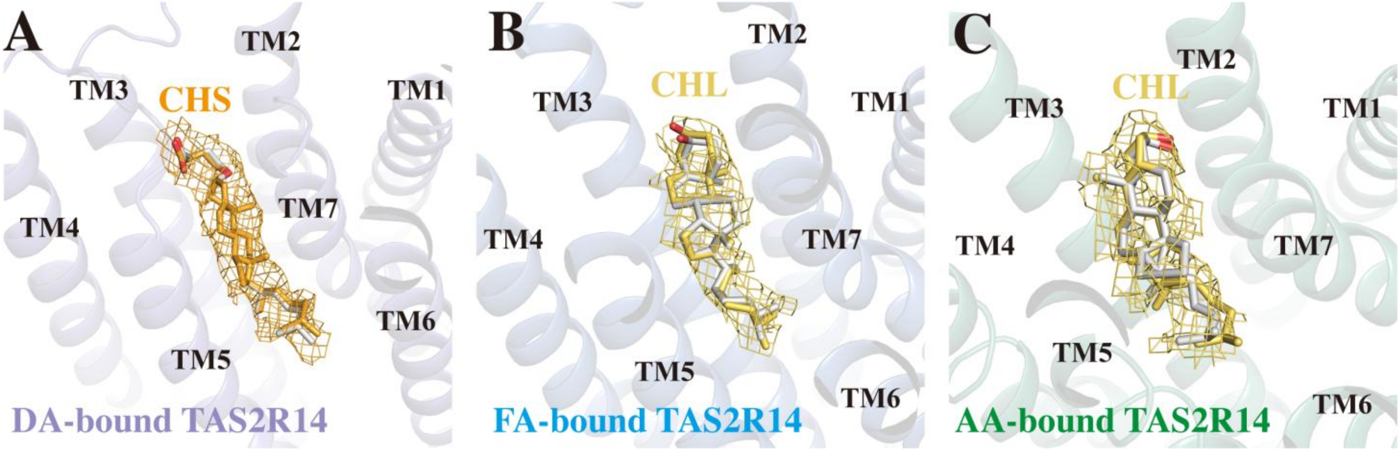


**Fig. S7. EMERALD docking results indicate CHS fits well with the density in DA-bound TAS2R14.** (A)The top-scoring pose for the CHS can recapitulate the modeled binding pose of the map. (B-C) EMERALD docking results of CHL poses in FA-bound (B) and AA-bound (C) TAS2R14 cryo-EM maps.

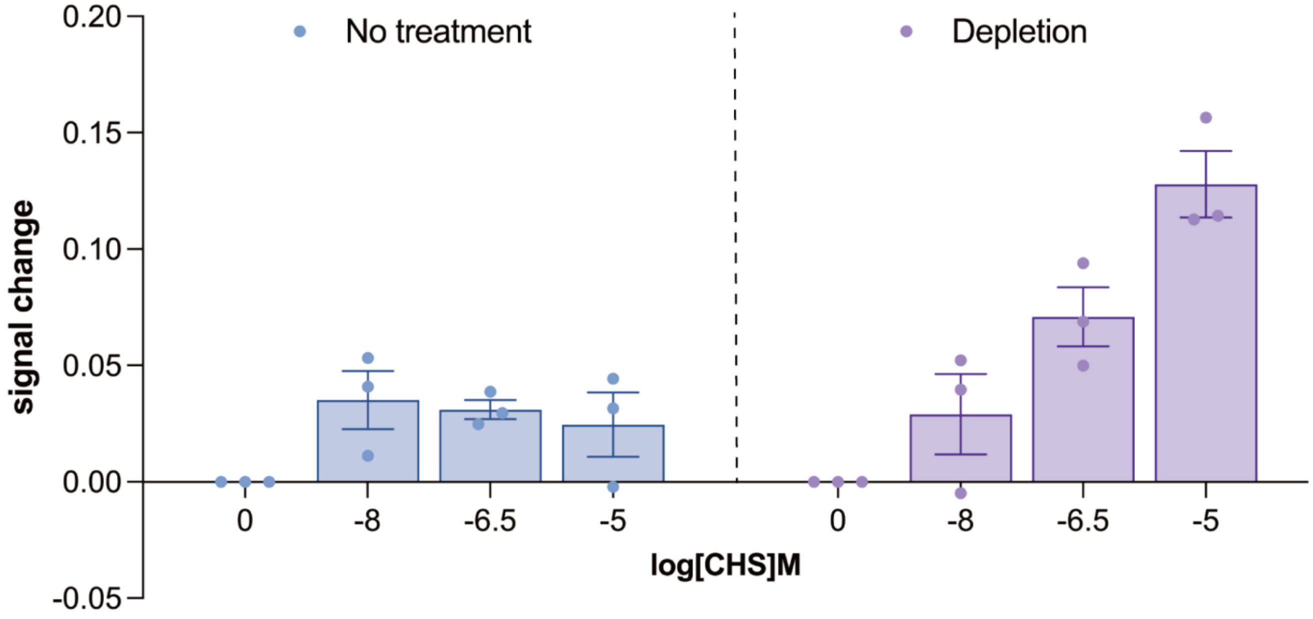


**Fig. S8. Calcium assays of CHS-stimulated TAS2R14 activation upon conditions of CHL depletion and no treatment.**

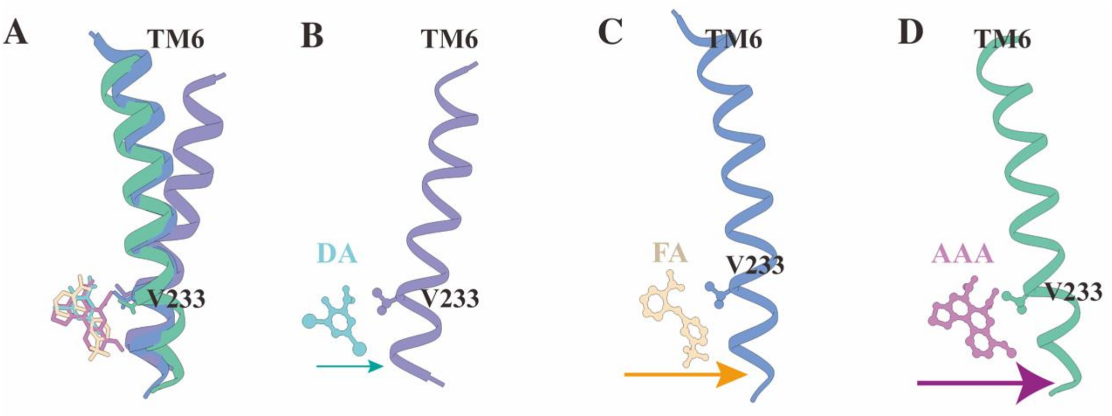


**Fig. S9. Comparison of TM6 in DA/FA/AA-bound TAS2R14 structures.** (**A**) Superimposition of TM6 in DA, FA and AA-bound TAS2R14 structures. (**B** to **D**) TM6 in TAS2R14 structures bound with DA (B), FA (C) and AA (D) with corresponding ligands.

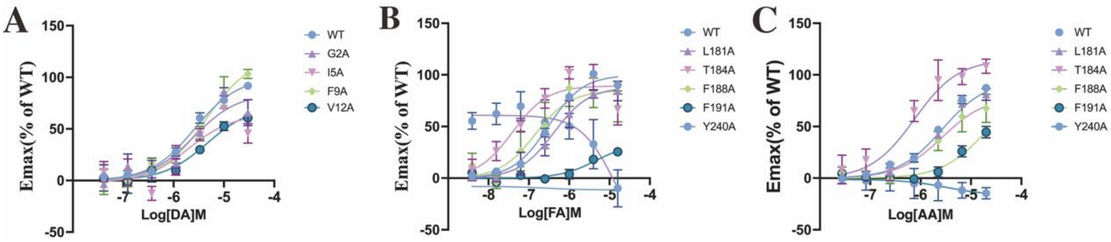


**Fig. S10. Dose responses curves of key residues mutations in TM1, TM5 and TM6 and receptor activation for DA, FA and AA in calcium mobilization assay. (A)** Dose–response curves of key residues mutations in TAS2R14 in response to stimulation with DA in calcium mobilization assay. Data are mean ± s.e.m. from 3 independent experiment (n = 3). **(B)** Dose–response curves of key residues mutations in TAS2R14 in response to stimulation with FA in calcium mobilization assay. Data are mean ± s.e.m. from 3 independent experiments (n = 3). **(C)** Dose–response curves of key residues mutations in TAS2R14 in response to stimulation with AA in calcium mobilization assay. Data are mean ± s.e.m. of $n = 3$ biological replicates.

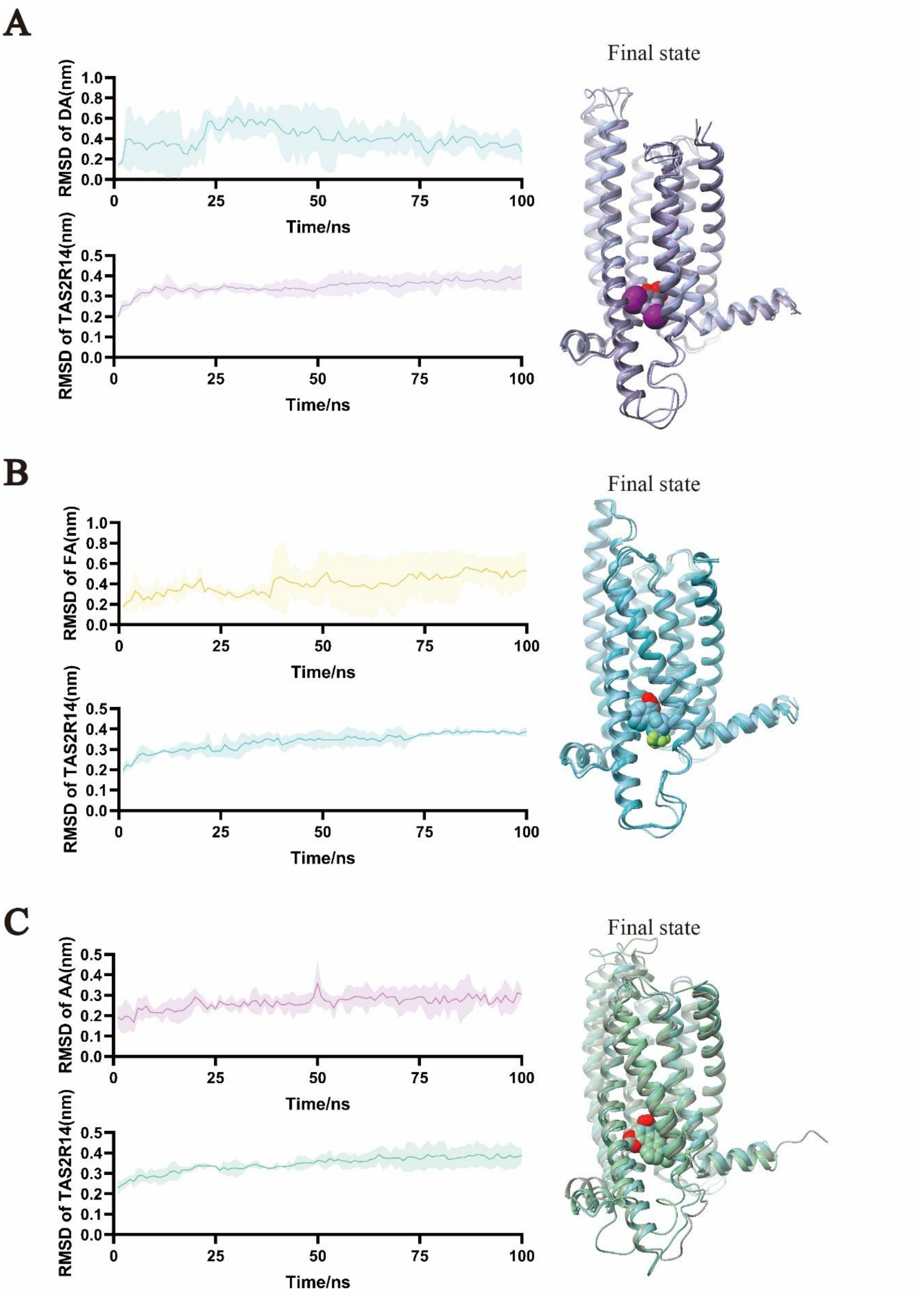


**Fig. S11. Molecular dynamic simulation of DA-TAS2R14 system (A), FA-TAS2R14 system (B), and AA-TAS2R14 system (C).**

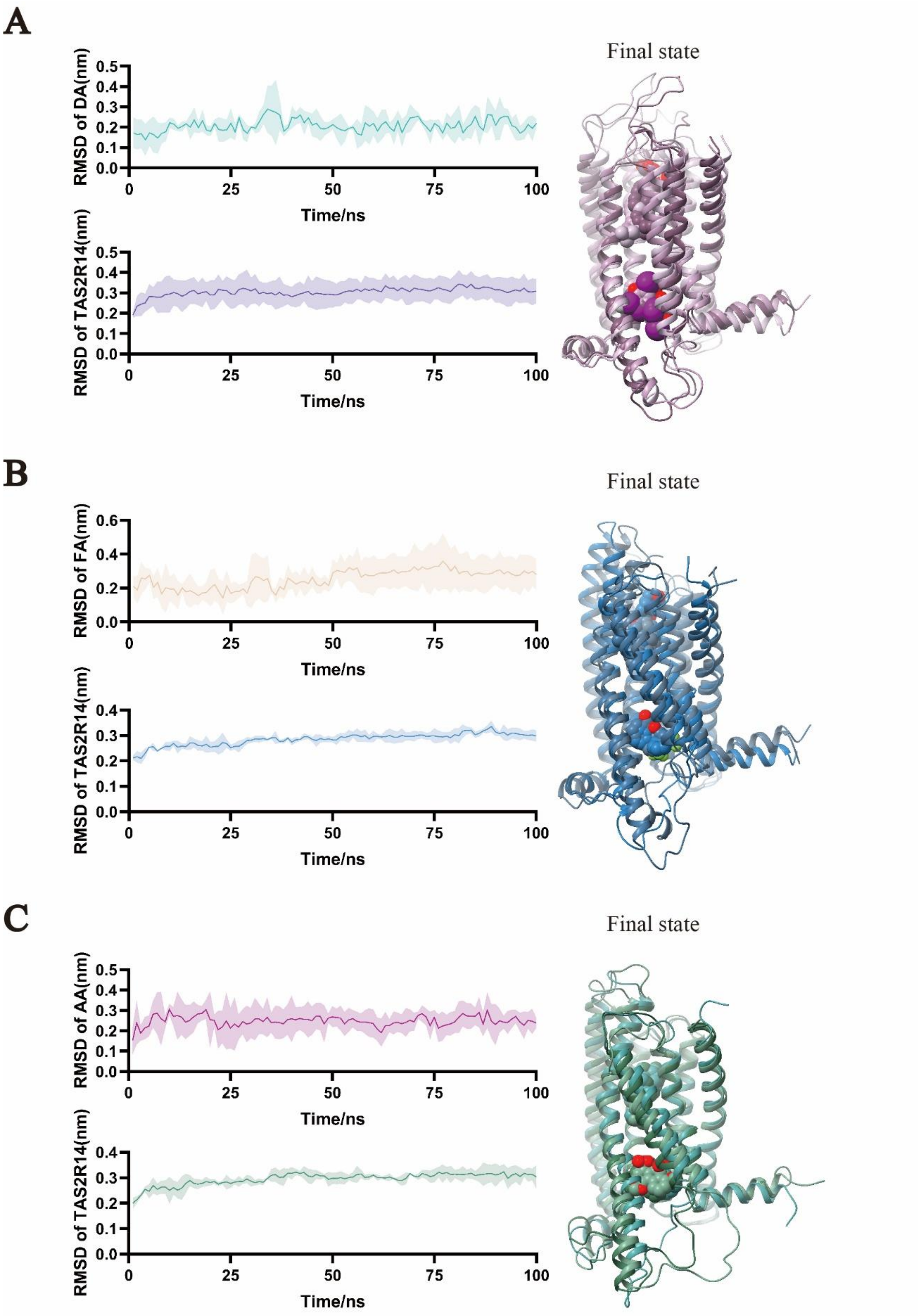


**Fig. S12. Molecular dynamic simulation of DA-CHS-TAS2R14 system (A), FA-CHL-TAS2R14 system (B), and AA-CHL-TAS2R14 system (C).**

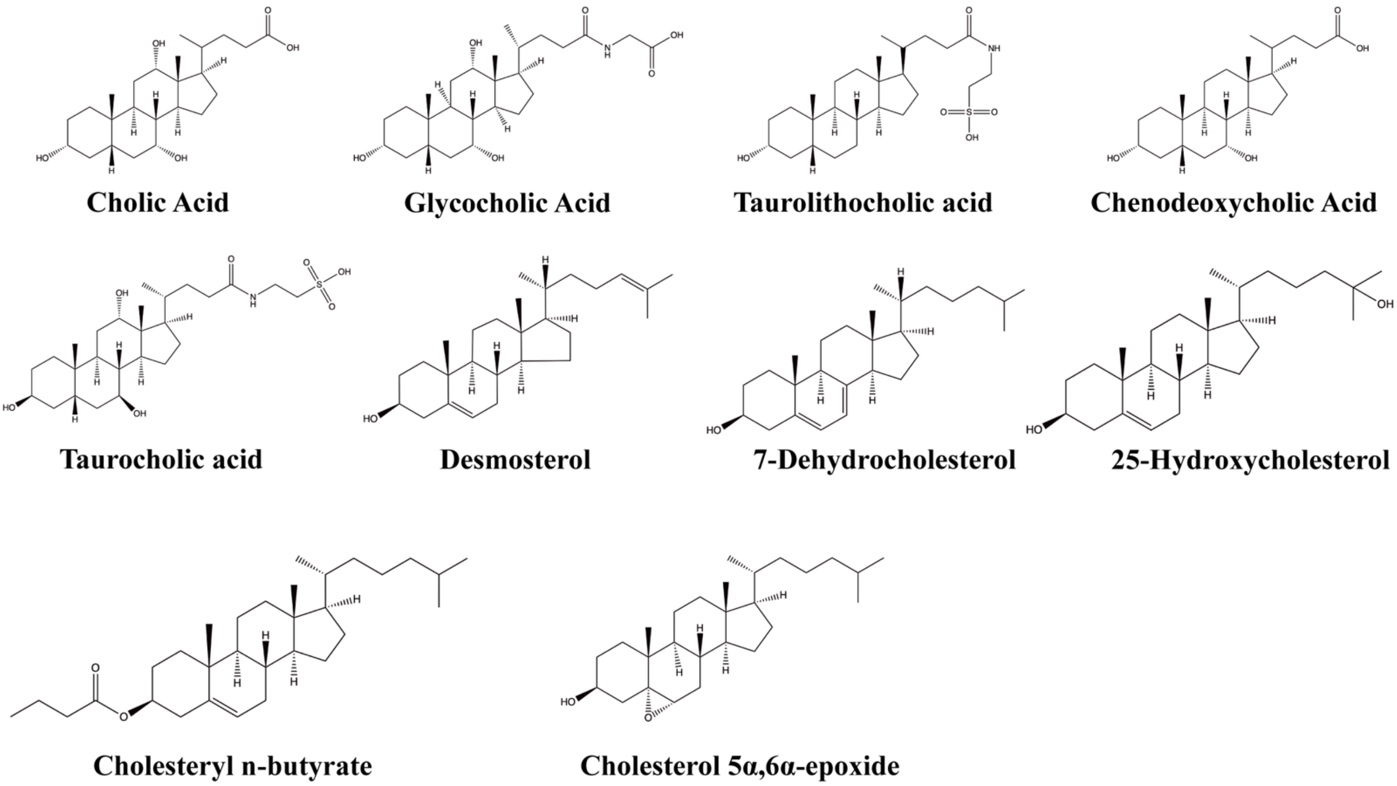


**Fig. S13. Distinct chemical scaffolds of cholesterol derivatives.**

**Table S1. Cryo-EM data collection, refinement and validation statistics**

| | 14-da complex | 14-fa complex | 14-aa complex |
|---|---|---|---|
| | ( ) | ( ) | ( ) |
| | ( ) | ( ) | ( ) |
| **Data collection and processing** | | | |
| Magnification | 105,000 | 105,000 | 105,000 |
| Voltage (kV) | 300 | 300 | 300 |
| Electron exposure ($e^{-}$/Å$^2$) | 54 | 54 | 54 |
| Defocus range (μm) | -1.0 ~ -1.5 | -1.0 ~ -1.5 | -1.0 ~ -1.5 |
| Pixel size (Å) | 0.851 | 0.851 | 0.851 |
| Symmetry imposed | C1 | C1 | C1 |
| Initial particle projections (no.) | 4,987,646 | 3,812,717 | 4,270,697 |
| Final particle projections (no.) | 1,385,801 | 1,296,323 | 1,166,258 |
| Map resolution (Å) | 2.44 | 2.39 | 2.69 |
| FSC threshold | 0.143 | 0.143 | 0.143 |
| Map resolution range (Å) | 1.86 ~ 21.37 | 1.86 ~ 9.41 | 2.27 ~ 10.50 |
| **Refinement** | | | |
| Initial model used | AlphaFold2 | XXXX | XXXX |
| Model resolution (Å) | 2.49 | 2.48 | 2.50 |
| FSC threshold | 0.5 | 0.5 | 0.5 |
| Map sharpening B factor (Å$^2$) | -110.6 | -106.0 | -129.2 |
| Model composition | | | |
| Non-hydrogen atoms | 8,959 | 9,111 | 8,873 |
| Protein residues | 1,149 | 1,158 | 1,129 |
| Ligand | 2 | 2 | 3 |
| *B*-factors (Å$^2$) | | | |
| Protein | 11.62/116.10/57.67 | 7.66/151.75/60.88 | 0.71/125.84/51.76 |
| Ligand | 121.55/121.55/121.55 | 84.64/84.64/84.64 | 86.58/91.24/88.91 |
| R.m.s. deviations | | | |
| Bond lengths (Å) | 0.006 | 0.002 | 0.003 |
| Bond angles (°) | 0.709 | 0.571 | 0.598 |
| Validation | | | |
| MolProbity score | 1.78 | 1.53 | 1.33 |
| Clashscore | 4.93 | 4.35 | 4.70 |
| Rotamer outliers (%) | 0.00 | 2.60 | 0.00 |
| Ramachandran plot | | | |

| | | | |
|---|---|---|---|
| Favored (%) | 96.64 | 97.99 | 97.57 |
| Allowed (%) | 3.36 | 2.01 | 2.43 |
| Disallowed (%) | 0.00 | 0.00 | 0.00 |

**Table S2. Diiodosalicylic acid(DA), Flufenamic Acid(FA), Aristolochic Acid A(AA)-induced intracellular calcium mobilization of wild-type (WT) and mutant TAS2R14.**

| Mutants | DA | | | | FA | | | | AA | | | | Expression (% WT) [a] |
|---|---|---|---|---|---|---|---|---|---|---|---|---|---|
| | $EC_{50}$ (μM) | pEC50±SEM[a] | Emax (% of WT)[a,b] | n[c] | $EC_{50}$ (μM) | pEC50±SEM[a] | Span (% of WT)[a,b] | n[c] | $EC_{50}$ (μM) | pEC50±SEM[a] | Span (% of WT)[a,b] | n[c] | |
| Wild type | 2.61 | 5.58±0.08 | 100±4 | 20 | 0.30 | 6.62±0.06 | 100±3 | 22 | 2.46 | 5.61±0.07 | 100±4 | 18 | 100 |
| Bril-TAS2R14 | 2.64 | 5.58±0.29 | 219±34** | 3 | 0.43 | 6.37±0.19 | 199±19*** | 3 | 2.11 | 5.67±0.23 | 160±21*** | 3 | 193±34*** |
| G2$^{1.28}$A | 2.51 | 5.60±0.24 | 84±11 | 3 | — | — | — | — | — | — | — | — | 94±44 |
| I5$^{1.31}$A | 1.99 | 5.70±0.37 | 66±13 | 3 | — | — | — | — | — | — | — | — | 65±16 |
| F9$^{1.35}$A | 4.44 | 5.35±0.17 | 119±12 | 3 | — | — | — | — | — | — | — | — | 53±3 |
| V12$^{1.38}$A | 4.65 | 5.33±0.26 | 72±11 | 3 | — | — | — | — | — | — | — | — | 130±43 |
| S25$^{1.51}$G | 3.53 | 5.45±0.31 | 115±20 | 3 | 0.51 | 6.29±0.18 | 96±9 | 3 | 4.52 | 5.34±0.21 | 101±15 | 3 | 94±14 |
| L103$^{3.46}$A | nd | nd | nd | 3 | nd | nd | nd | 3 | nd | nd | nd | 3 | 81±9 |
| G104$^{3.47}$A | 7.50 | 5.13±0.23 | 78±12 | 3 | 1.43 | 5.84±0.12 | 92±6 | 3 | 4.60 | 5.34±0.16 | 106±12 | 3 | 57±4 |
| G104$^{3.47}$L | nd | nd | nd | 3 | nd | nd | nd | 3 | nd | nd | nd | 3 | 78±5 |
| G104$^{3.47}$S | 1.94 | 5.72±0.27 | 81±12 | 3 | 0.36 | 6.44±0.13 | 133±8 | 3 | nd | nd | nd | 3 | 110±9 |
| Y107$^{3.50}$A | nd | nd | nd | 3 | nd | nd | nd | 3 | nd | nd | nd | 3 | 90±13 |
| Y107$^{3.50}$F | nd | nd | nd | 3 | nd | nd | nd | 3 | nd | nd | 46±32*** | 3 | 109±15 |
| F108$^{3.51}$A | nd | nd | nd | 3 | nd | nd | nd | 3 | nd | nd | nd | 3 | 67±5 |

| | | | | | | | | | | | | | |
|---|---|---|---|---|---|---|---|---|---|---|---|---|---|
| L181$^{5.41}$A | 1.84 | 5.74±0.38 | 122±24 | 3 | 0.42 | 6.38±0.19 | 88±8 | 3 | 2.70 | 5.57±0.07 | 95±4 | 3 | 82±19 |
| T184$^{5.44}$A | — | — | — | — | 0.04 | 7.40±0.29 | 89±15 | 3 | 0.76 | 6.12±0.22 | 115±13 | 3 | 61±12 |
| F188$^{5.48}$A | 3.66 | 5.44±0.38 | 108±24 | 3 | 0.14 | 6.84±0.36 | 85±16 | 3 | 2.66 | 5.58±0.25 | 79±12 | 3 | 143±24 |
| F191$^{5.51}$A | — | — | — | — | 4.44 | 5.35±0.27** | 33±7*** | 3 | nd | nd | 68±14** | 3 | 138±13 |
| S194$^{5.54}$A | nd | nd | nd | 3 | nd | nd | nd | 3 | nd | nd | nd | 3 | 73±11 |
| M197$^{5.57}$A | nd | nd | nd | 3 | 0.68 | 6.17±0.13 | 64±4 | 3 | nd | nd | 31±4*** | 3 | 69±6 |
| F198$^{5.58}$A | 1.90 | 5.72±0.12 | 167±11* | 3 | 0.52 | 6.30±0.29 | 74±11 | 3 | nd | nd | nd | 3 | 42±5 |
| L201$^{5.61}$A | nd | nd | nd | 3 | 1.18 | 5.93±0.31 | 49±8** | 3 | nd | nd | 39±18*** | 3 | 97±13 |
| I202$^{5.62}$A | nd | nd | nd | 3 | 0.92 | 6.04±0.34 | 49±9** | 3 | nd | nd | 32±1*** | 3 | 97±18 |
| M205$^{5.65}$L | 10.64 | 4.97±0.29 | 116±30 | 3 | 0.37 | 6.43±0.18 | 72±7 | 3 | 2.11 | 5.68±0.17 | 77±7 | 3 | 91±12 |
| A226$^{6.31}$W | nd | nd | nd | 3 | 0.07 | 6.98±0.55 | 24±7*** | 3 | 2.14 | 5.67±0.32 | 44±8*** | 3 | 124±11 |
| A226$^{6.31}$L | nd | nd | nd | 3 | 0.20 | 6.70±0.30 | 51±8*** | 3 | nd | nd | 33±6*** | 3 | 104±23 |
| G229$^{6.34}$F | nd | nd | nd | 3 | nd | nd | nd | 3 | nd | nd | nd | 3 | 87±8 |
| V230$^{6.35}$A | nd | nd | nd | 3 | 0.46 | 6.33±0.22 | 60±6 | 3 | nd | nd | 52±11* | 3 | 109±22 |
| V233$^{6.38}$A | nd | nd | nd | 3 | nd | nd | nd | 3 | nd | nd | nd | 3 | 62±7 |
| F237$^{6.42}$A | nd | nd | nd | 3 | nd | nd | nd | 3 | nd | nd | nd | 3 | 70±9 |
| F237$^{6.42}$Y | 4.42 | 5.35±0.36 | 180±39 | 3 | 0.08 | 7.07±0.17 | 162±14** | 3 | 5.60 | 5.25±0.25 | 85±16 | 3 | 58±8 |
| Y240$^{6.45}$A | — | — | — | — | nd | nd | nd | 3 | nd | nd | nd | 3 | 63±8 |

| | | | | | | | | | | | | | |
|---|---|---|---|---|---|---|---|---|---|---|---|---|---|
| F243$^{6.48}$A | 6.42 | 5.19±0.29 | 95±19 | 3 | 0.24 | 6.61±0.29 | 116±17 | 3 | 1.97 | 5.71±0.26 | 74±11 | 3 | 109±17 |
| S244$^{6.49}$A | 1.75 | 5.76±0.21 | 123±13 | 3 | 0.18 | 6.74±0.29 | 183±26*** | 3 | 1.83 | 5.74±0.16 | 113±10 | 3 | 63±5 |
| F247$^{6.52}$A | nd | nd | nd | 3 | nd | nd | nd | 3 | nd | nd | nd | 3 | 123±31 |
| F248$^{6.53}$A | 3.61 | 5.44±0.25 | 162±24 | 3 | 0.16 | 6.79±0.38 | 167±32** | 3 | 3.47 | 5.46±0.17 | 73±8 | 3 | 67±17 |
| S250$^{6.55}$A | 1.29 | 5.89±0.27 | 268±37*** | 3 | 0.12 | 6.92±0.35 | 91±16 | 3 | 2.70 | 5.57±0.22 | 126±16 | 3 | 117±18 |
| W252$^{6.57}$A | 2.63 | 5.58±0.32 | 112±18 | 3 | 0.21 | 6.67±0.28 | 93±13 | 3 | 1.18 | 5.93±0.13 | 62±4 | 3 | 70±5 |
| H276$^{7.49}$A | nd | nd | nd | 3 | nd | nd | nd | 3 | nd | nd | nd | 3 | 90±7 |
| V279$^{7.52}$A | nd | nd | nd | 3 | 0.21 | 6.68±0.39 | 52±10 | 3 | nd | nd | 21±3*** | 3 | 103±5 |
| L280$^{7.53}$A | nd | nd | nd | 3 | nd | nd | nd | 3 | nd | nd | nd | 3 | 82±6 |
| G283$^{7.56}$T | 1.43 | 5.84±0.16 | 153±12 | 3 | 0.21 | 6.67±0.35 | 74±13 | 3 | 0.19 | 6.73±0.33*** | 74±14 | 3 | 71±10 |

[a]Data are shown as mean±SEM from at least three independent experiments performed in technical triplicate. *P<0.05; **P<0.01; ***P<0.001 by one-way ANOVA followed by Dunnett's post-test, compared with the response of the WT. nd (not determined) refers to data where a robust concentration response curve could not be established within the concentration range tested.

[b] The maximal response $E_{max}$ values were calculated from the concentration-response curves.

[c]Sample size; the number of independent experiments performed in technical triplicate.

**Table S3. Diiodosalicylic acid(DA), Flufenamic Acid(FA), Aristolochic Acid A(AA)-induced intracellular calcium mobilization of TAS2R14 with different treatments.**

| Treatment | DA | | | | FA | | | | AA | | | |
|---|---|---|---|---|---|---|---|---|---|---|---|---|
| | $EC_{50}$ (μM) | pEC50±SEM[a] | Span (% of no treatment )[a,b] | n[c] | $EC_{50}$ (μM) | pEC50±SEM[a] | Span (% of no treatment)[a,b] | n[c] | $EC_{50}$ (μM) | pEC50±SEM[a] | Span (% of no treatment)[a,b] | n[c] |
| No treatment | 2.54 | 5.60±0.15 | 100±8 | 7 | 0.20 | 6.70±0.08 | 100±4 | 10 | 2.26 | 5.65±0.18 | 100±10 | 3 |
| ΔCHL | 3.03 | 5.52±0.22 | 140±24** | 5 | 0.36 | 6.45±0.23 | 83±10 | 8 | 8.22 | 5.08±0.25 | 139±30 | 3 |
| +CHS | 1.09 | 5.96±0.30 | 119±18 | 4 | 0.32 | 6.45±0.20 | 105±10 | 4 | 1.39 | 5.86±0.17 | 154±14 | 3 |
| ΔCHL+CHS | 0.52 | 6.29±0.26*** | 120±18 | 5 | 0.41 | 6.38±0.17 | 111±9 | 8 | 5.95 | 5.23±0.16 | 135±17 | 3 |

[a]Data are shown as mean±SEM from at least three independent experiments performed in technical triplicate. *P<0.05; **P<0.01; ***P<0.001 by one-way ANOVA followed by Dunnett's post-test, compared with the response of no treatment.

[b] The maximal response $E_{max}$ values were calculated from the concentration-response curves.

[c]Sample size; the number of independent experiments performed in technical triplicate.

**Table S4. Diiodosalicylic acid (DA), Flufenamic Acid (FA), Aristolochic Acid (AA) -induced intracellular calcium mobilization of wild-type (WT) TAS2R14.**

| Ligands | $EC_{50}$ (μM) | pEC50±SEM[a] | Emax (% of FA)[a,b] | n[c] |
|---|---|---|---|---|
| DA | 2.15 | 5.57±0.11 | 53±3** | 6 |
| FA | 0.30 | 6.20±0.10 | 100±5 | 6 |
| AA | 2.34 | 5.63±0.12 | 136±9* | 6 |

[a]Data are shown as mean±SEM from at least three independent experiments performed in technical triplicate. *P<0.05; **P<0.01; ***P<0.001 by one-way ANOVA followed by Dunnett's post-test, compared with the response of the FA.

[b] The maximal response $E_{max}$ values were calculated from the concentration-response curves.

[c]Sample size; the number of independent experiments performed in technical triplicate.
